# A Novel Engineered U7 Small Nuclear RNA Scaffold Greatly Increases *in vitro* and *in vivo* ADAR-Mediated Programmable RNA Base Editing

**DOI:** 10.1101/2024.09.29.615721

**Authors:** Susan M. Byrne, Stephen M. Burleigh, Robert Fragoza, Yue Jiang, Yiannis Savva, Ricky Pabon, Joseph Rainaldi, Andrew Portell, Prashant Mali, Adrian W. Briggs

## Abstract

Custom RNA base editing using the endogenous human Adenosine Deaminase Acting on RNA (ADAR) enzyme presents a promising approach for precision therapeutics, alleviating concerns of permanent DNA damage or immunogenicity from^1^ foreign bacterial proteins such as CRISPR/Cas. ADAR can be directed to act on therapeutic RNA targets by antisense guide RNAs (gRNAs) that create a substrate for ADAR’s adenosine-to-inosine (effectively A-to-G) deamination activity. Delivery of gRNAs via a DNA expression construct provided by Adeno-Associated Virus (AAV) might allow life-long duration of the therapy. However, a major challenge for RNA editing using gene-encoded gRNAs and endogenous levels of ADAR is achieving sufficient gRNA activity inside cells, especially in therapeutic situations where AAV delivery may provide as low as one viral genome per cell. Here we show that embedding antisense gRNAs into a U7 small nuclear RNA (snRNA) framework and adding hnRNP A1 binding domains greatly increases the efficiency of custom RNA editing. This increased editing efficiency allows for detectable RNA editing from a single genomic insertion of gRNA construct per cell, which enabled a pooled library screen of 750+ gRNA variations to further improve the SmOPT U7 hairpin system. The screen revealed critical residues responsible for RNA editing and generated new SmOPT and U7 hairpin variants that further boosted RNA editing. The final design, combined with an improved synthetic U7 promoter, resulted in up to 76% targeted editing with a single integrated copy of construct per cell, representing a 10- to 100-fold increase over existing circular gRNA approaches. Using systemic *in vivo* AAV delivery, we achieved an unprecedented 75% RNA editing in the total brain of a mouse model of Hurler syndrome. Our novel SmOPT U7 system also improved published antisense oligos for DMD exon skipping, currently in clinical trials, by up to 25-fold in differentiated myoblasts, and therefore represents a universal scaffold for ADAR-based RNA editing as well as other antisense RNA therapies.

---

Editing nucleic acid sequences *in vivo* holds massive potential for precisely treating genetic diseases and beyond. While CRISPR/Cas9 nucleases and DNA base editors have revolutionized gene and cell therapies, drawbacks remain, such as the potential for DNA damage, permanent off-targets, or germline heritability, as well as delivery challenges and immunogenicity from foreign proteins. In contrast to DNA editing with bacterial enzymes, custom RNA editing using endogenous human ADAR enzymes may eliminate these concerns^1,2^. ADAR binds double-stranded RNA (dsRNA) structures and deaminates adenosines to inosine, which the cellular translation machinery interprets as guanosine. Natural ADAR deamination plays many important functions: besides editing mRNA coding sequences, it regulates RNA splice sites, modulates RNA interference, and shields endogenous dsRNA including *Alu* repeat sequences from immune sensors of viral dsRNA^3,4^. For therapeutic purposes, ADAR can be recruited to additional adenosines using a custom antisense gRNA that anneals to the target RNA sequence to form the dsRNA^5^.

For decades, it has been known that custom antisense RNA sequences for therapeutic exon skipping can be substantially enhanced using the framework of a natural U7 small nuclear RNA (snRNA)^6–8^. The human U7 snRNA normally binds histone pre-mRNA through a complementary 20nt Histone Downstream Element (HDE) and directs endonucleocytic cleavage of the histone pre-mRNA, in a process that also depends on the Sm-protein binding region and the downstream hairpin of the U7 snRNA. By replacing the HDE with a 20-30 nt custom antisense sequence, optimizing the Sm-binding domain (SmOPT), and driving expression of the construct by the U7 promoter, it was found that exon skipping of a therapeutic target RNA could be substantially increased over the antisense sequence alone^9,10^. Associated studies revealed that the assembled Sm protein complex protects the short antisense RNAs from degradation, and the standard 5’ RNA cap becomes hypermethylated to a 2,2,7-trimethyl guanosine (3mG) cap for nuclear localization^11^. Importantly, the engineered antisense SmOPT U7 RNAs did not adversely affect natural U7 snRNA histone processing, the broader transcriptome, or display other toxicities^12,13^. This strategy has now been tested against a wide variety of diseases^14–19^; notably, an ongoing clinical trial involving skipping of a duplicated DMD exon 2 showed a sustained benefit in patients up to 18 months post treatment ^20–22^.

Natural ADAR editing and splicing frequently occur co-transcriptionally in the nucleus^4,23^. Therefore in this study, we hypothesized that the same benefits provided to exon skipping by the U7 snRNA framework might also apply to ADAR-based targeted RNA editing. Here we show that the SmOPT U7 hairpin dramatically increases the potency of 60-100 nt antisense RNAs for ADAR-mediated RNA editing. The greatly increased efficiency of ADAR editing using the SmOPT U7 system removes the need for ADAR enzyme overexpression and allows detectable editing from a single DNA copy of the gRNA expression construct. This breakthrough allowed us to perform a pooled library screen exploring the effect of mutations along the original SmOPT and U7 hairpin framework. This screen shed light on potential mechanisms by identifying critical residues required for RNA editing, as well as revealing improved SmOPT U7 hairpin variants that further boosted RNA editing. In particular, an improved triple-variant SmOPT U7 hairpin, with an engineered enhanced U7 promoter, resulted in 76% targeted RNA editing from a single integrated DNA copy of the gRNA construct, and 75% editing in total, unsorted mouse brain from a systemic AAV PHP.eB injection. Furthermore, when the new SmOPT U7 hairpin variants and promoters were coupled to the previously published antisense constructs for DMD, exon skipping increased up to 25-fold in differentiated myoblasts.

## Results

### Expression of ADAR gRNAs in a SmOPT U7 format boosts RNA editing

Guide RNAs for custom ADAR editing have traditionally been driven by the U6 snRNA promoter, which is also widely used to express short hairpin RNAs and CRISPR gRNAs. We previously reported^1,24^ that a U6-driven 100 nt antisense RNA possessing an A-C mismatch in the center (denoted 100.50) enables editing of the RAB7A transcript by endogenous ADAR, although editing efficiencies are higher upon ADAR overexpression. Here, we found that when this antisense gRNA was coupled to an “Optimal” Sm-binding (SmOPT) sequence [AAUUUUUGGAG]^9^ and a human U7 (hU7) or mouse U7 (mU7) snRNA hairpin^25,26^, and expressed via a U7 promoter, RNA editing with endogenous ADAR was greatly increased to levels comparable to those with ADAR overexpression (Fig. 1a), thus removing the need for additional enzyme. Excitingly, similar patterns of robust RNA editing were seen for antisense gRNAs targeting FANCC 3’UTR, SMAD4 coding sequence, and SOD1 translation initiation site (TIS) (Fig. 1a), as well as splice acceptor adenosines for DMD exons 71 and 74 (Suppl. Fig. 1). While the RAB7A, SOD1, and DMD antisense sequences comprised a simple A-C mismatch, the FANCC and SMAD4 antisense sequences also contained mismatch loops at the −5 and +30 positions, previously shown to improve editing specificity^1^. We further confirmed that both elements of the U7 snRNA framework were necessary – *i*.*e*. neither the SmOPT U7 hairpin nor the promoter alone increased RNA editing (Fig. 1a). Further testing showed that the SmOPT sequence was required; substituting it with the wild type U7 or U1 Sm binding domains sharply reduced activity (Suppl. Fig. 2).

**Fig. 1.**
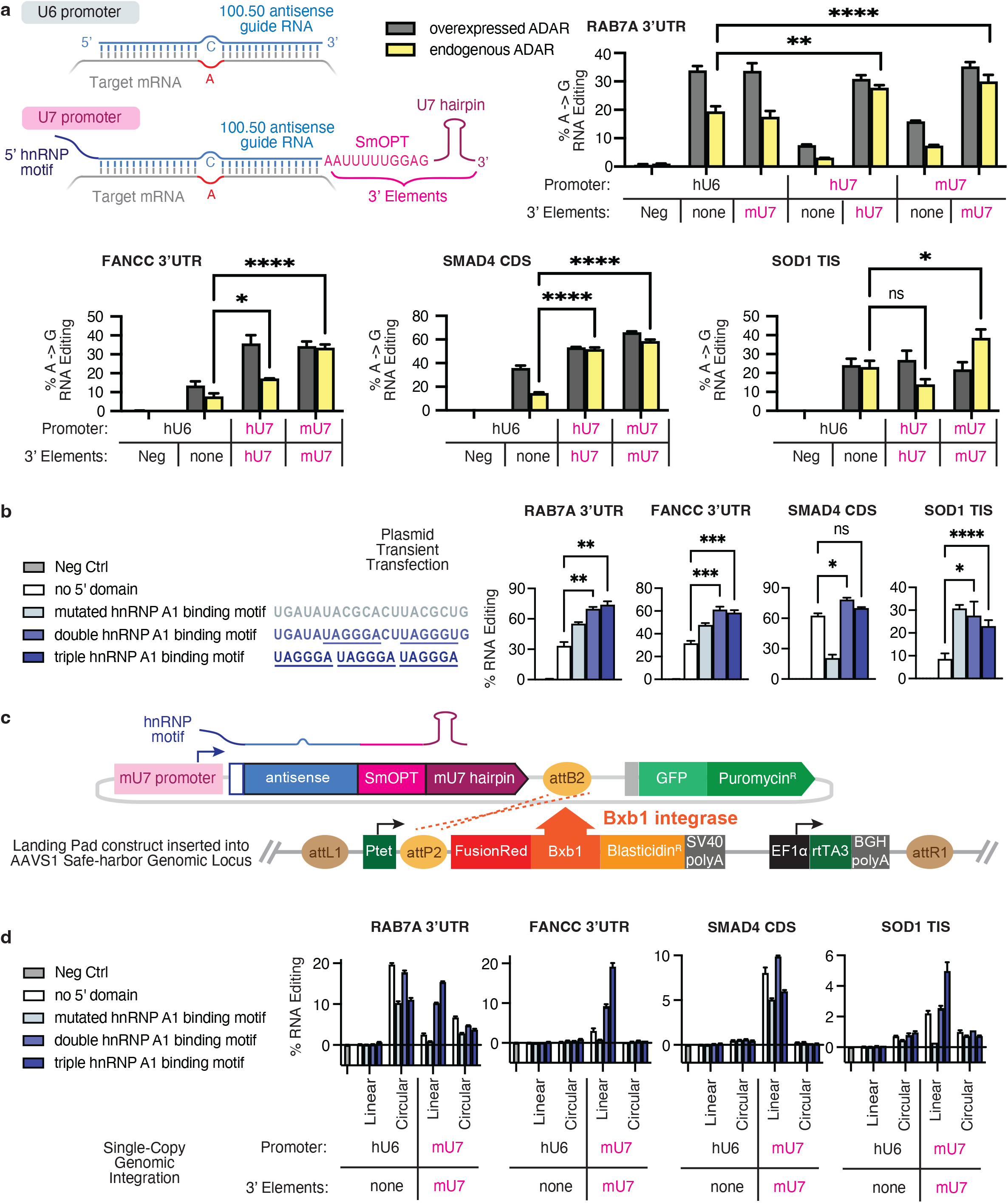
Adding a U7 promoter and SmOPT U7 hairpin to the antisense gRNA improves RNA editing. **a**, 100 nt antisense guide RNAs targeting adenosines in the RAB7A 3’UTR, FANCC 3’UTR, SMAD4 coding sequence, or SOD1 start codon (TIS) were expressed using the hU6, hU7, or mU7 promoters, either alone or possessing an additional SmOPT sequence and human or mouse U7 hairpin on the 3’end. Plasmids also contained the CMV promoter to overexpress ADAR2 (gray bars) or GFP (endogenous ADAR, yellow bars). RNA was measured from HEK293T cells 2 days post transfection. **b**, Sequences containing two or three copies of the hnRNP A1 binding motif (UAGGGW, underlined) were added onto the 5’ end of antisense guide RNAs expressed using a mouse U7 promoter and possessing a SmOPT mU7 hairpin. RNA was measured from HEK293T cells with endogenous ADAR levels 2 days after plasmid transfection. **c**, Landing Pad system for testing guide RNAs at a single copy per cell. A Landing Pad construct allowing doxycycline-inducible expression of FusionRed, BxB1 integrase, and a blasticidin-resistance marker was integrated into a single AAVS1 safe harbor genomic locus in HEK293T cells. When the resulting “Landing Pad” cell line was transfected with plasmids expressing a guide RNA and selectable marker, BxB1 integrase recombines the attB2 and attP2 sites to incorporate exactly one guide RNA construct into a single site in the genome. **d**, Antisense guide RNAs possessing various 5’ hnRNP A1 domains were constructed using the hU6 promoter with no 3’hairpin or the mU7 promoter with a SmOPT mU7 hairpin as described above (Linear) or containing additional circularizing ribozyme sequences flanking the guide RNA (Circular). Constructs were transfected into HEK293T Landing Pad cells; RNA from successfully integrated single-copy gRNA constructs was measured 13 days after the initial plasmid transfection. Negative controls were measured from samples where the gRNA targeted a different gene. Significance was calculated by Mixed-effects analysis using Dunnett’s multiple comparisons test.

**Fig. 2.**
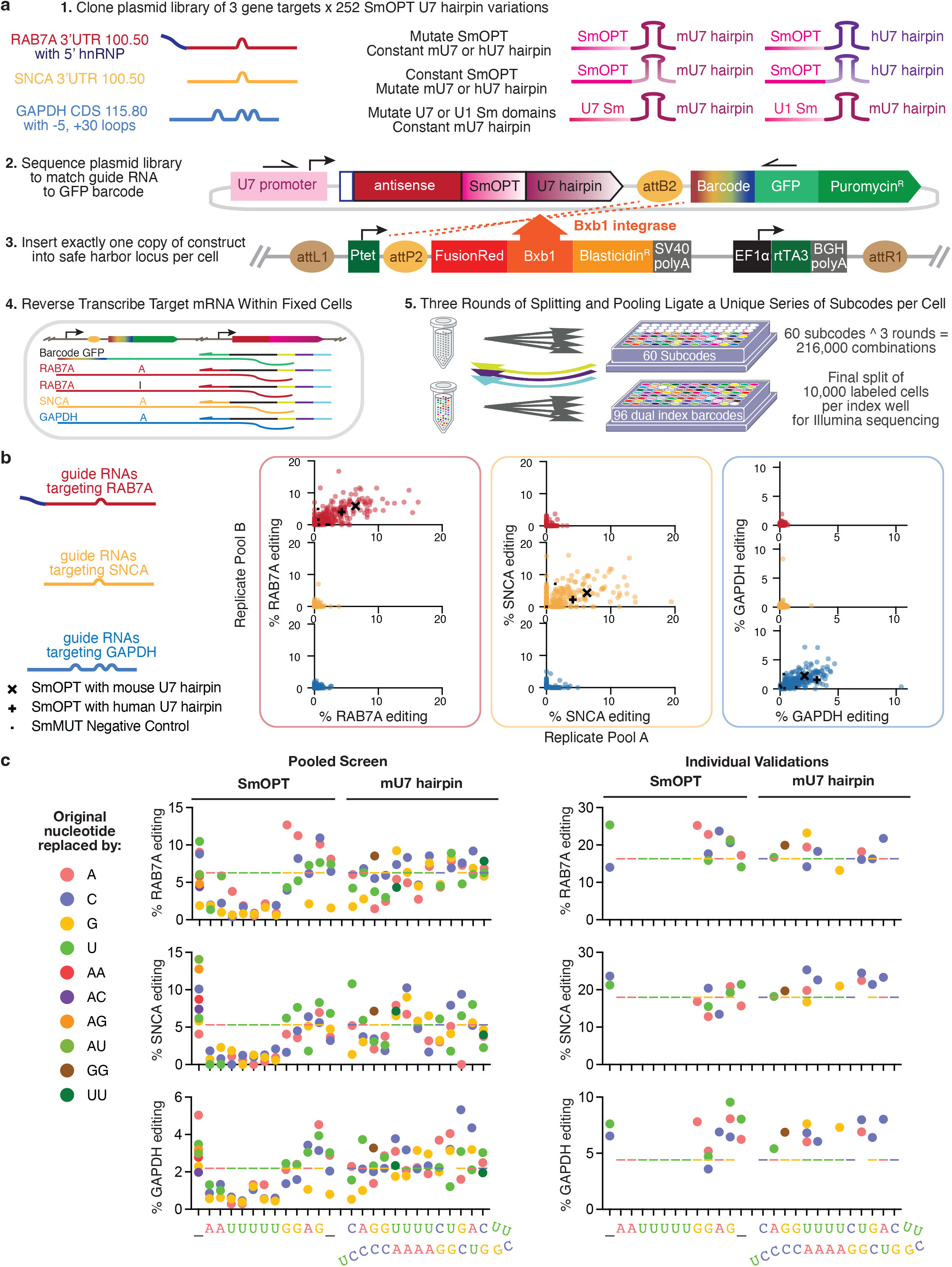
Pooled screen of SmOPT and U7 hairpin variants using split-pool single cell barcoding. **a**, Mutagenesis screen workflow. 1) Three antisense guide RNAs targeting the RAB7A 3’UTR, SNCA 3’UTR, or GAPDH CDS were each cloned onto 252 mutagenesis variants along the Sm binding or U7 hairpin sequences. Duplicate plasmid libraries were cloned for each target gene, yielding six pools total. 2) Plasmid libraries were sequenced to match the guide RNA with a 14N random “GuideID” barcode located in the GFP 5’UTR. 3) Plasmid libraries were transfected into the Landing Pad 293T cell line; after successful integration, 10M cells, each containing a single copy of construct, were pooled, fixed, and permeabilized. 4) Four custom primers reverse transcribed GFP, RAB7A, SNCA, or GAPDH transcripts within the fixed cell. Each RT primer possessed an additional 10N umi and an anchor sequence, which was annealed to a Splint scaffold. 5) Cells were split equally across 60 different wells, each containing a different 7mer subcode. After annealing to the Splint scaffold, subcodes were ligated onto the reverse transcribed cDNA inside the fixed cells. Cells were then washed to remove unligated subcodes, pooled, and split again across 60 new wells. Three rounds of split-pool ligation across 60 subcode wells results in 60^3 or 216,000 possible subcode combinations. Finally, 10,000 labeled cells were aliquoted into each well of a PCR plate, lysed, and amplified with dual index primers for Illumina sequencing. **b**, For each of the 750+ guide RNAs, RNA editing was measured for all three transcripts. Editing results comparing the duplicate plasmid libraries (Replicate Pools A versus B) are shown. Guide RNAs targeting RAB7A, SNCA, or GAPDH are shown in red, yellow, or blue, respectively. For each gene target, positive and negative controls for editing are marked by black symbols. **c**, For each gene target, the effect on RNA editing of single nucleotide substitutions across the SmOPT sequence or paired substitutions across the mU7 hairpin is shown for the corresponding antisense guide RNA. Underscores indicate locations where extra nucleotides were inserted. In some cases, two nucleotides were inserted. The multicolor dashed line marks the percent editing from the original SmOPT mU7 hairpin construct. Left panels show RNA editing rates from the pooled screen; right panels show RNA editing rates when desired mutations were individually validated at a single copy in the Landing Pad cell line.

### The hnRNP A1 binding site further enhances editing

To further enhance editing rates, we next explored coupling of additional domains for recruiting RNA binding proteins. In particular, we focused on Heterogenous nuclear ribonucleoprotein A1 (hnRNP A1), one of the most abundant nuclear proteins which has multiple functions in RNA processing, including: splicing modulation, transcriptional regulation, and intracellular localization^27^. Recruitment of hnRNP A1 has been observed to enhance antisense oligos for exon skipping^28,29^. When we attached hnRNP A1 binding motifs onto the 5’end of our antisense gRNAs, opposite the SmOPT and U7 hairpin, we observed an increase in RNA editing up to two-fold (Fig. 1b), although the effect varied depending on the antisense length and target.

### SmOPT-U7 gRNAs enable detectable editing from a single DNA copy of construct per cell

While a transient transfection may deliver hundreds of copies of plasmid to each cell, current levels of AAV gene delivery *in vivo* are often limited to one to few copies per cell. To compare our gRNAs in a stringent, controlled manner, we used the BxB1 integrase, which recombines attP and attB sites in a directed fashion, to insert exactly one copy of plasmid construct into the AAVS1 safe harbor genomic locus of 293T cells^30,31^. To maximize BxB1 integration efficiency, a “Landing Pad” cell line was created to provide doxycycline-inducible expression of the BxB1 integrase (Fig. 1c). Upon transfection, an attB-containing plasmid becomes integrated into the attP site within the Landing Pad construct. Afterwards, Landing Pad cells containing successful single-copy integrants are enriched by antibiotic selection, and editing can be measured in the stably transfected cell lines. With the U7 snRNA framework, we observed detectable RNA editing from a single copy of gRNA under endogenous ADAR levels (Fig. 1d). As before, adding a 5’ hnRNP A1 binding motif often further increased RNA editing. In contrast to SmOPT U7 gRNAs, antisense gRNAs expressed by the U6 promoter and lacking the SmOPT U7 hairpin showed no editing at this lower dose. Previous publications have shown that circular ADAR-recruiting gRNAs have improved durability and RNA editing activity^1,2^. When the antisense gRNA is expressed between flanking Twister ribozyme sequences, the ribozymes undergo autocatalytic cleavage and the RNA ligase RtcB inside the cell joins the RNA ends together to form a circle^32^. When antisense gRNAs with a SmOPT U7 hairpin and 5’ hnRNP A1 binding motif were flanked by Twister ribozymes, circular gRNAs were formed (data not shown), yet RNA editing was diminished compared to gRNAs with a linear U7 snRNA framework (Fig. 1d).

### High throughput split-pool sequencing enabled pooled screening of SmOPT-U7 variants

One benefit afforded by detectable RNA editing with single-copy DNA is the ability to perform a high throughput pooled screen on gRNAs. Using a single cell sequencing method, each cell’s RNA editing can be matched to the individual gRNA present within that cell, even though the gRNA-target RNA interaction occurs *in trans*. In contrast, previous pooled ADAR gRNA screens have only sequenced *in-cis* hairpin structures, where the target RNA is physically connected to the gRNA^33,34^. First, we needed to customize a system to link endogenous mRNA editing with gRNA identity in a single-cell context, since most methods of single cell RNA sequencing (scRNA-seq) only capture the 5’ or 3’ ends of transcripts and not the regions within a transcript targeted for editing. A recent method of split-pool single-cell barcoding used *in situ* reverse transcription to detect specific transcripts within a cell, assisted by a Splint scaffold to organize the cell-specific ligated barcodes^35^. We adapted this method to sequence longer (>100 nt) stretches with sufficient transcript capture efficiency and resolution to measure RNA editing at a specific target site. Protocol improvements included: moving the universal molecular identifier (umi) onto the reverse transcription (RT) primer for better fidelity; an alternative RT enzyme active at hotter temperatures; reducing four rounds of subcode ligation to three, with Illumina indexing supplying an additional level of barcode diversity; and an improved computational analysis pipeline.

Using this method, we performed a mutagenesis screen across the SmOPT and U7 hairpin sequences. Since it was first developed in 1993, the SmOPT sequence has been used with a U7 hairpin to target a wide range of diseases. However, while many groups have exchanged the antisense oligo portion of the short RNA, the SmOPT U7 hairpin sequence has remained unchanged^6^. Mutagenesis screens on natural U7 snRNA have uncovered some required nucleotides but also some flexibility within the Sm-binding site and U7 hairpin^36^. To discover which residues were essential for RNA editing, and possible improved variants, we generated a library of 252 mutations along the Sm binding or U7 hairpin sequences, comprising: single base substitutions across the SmOPT sequence with a constant mU7 or hU7 hairpin; a constant SmOPT sequence with paired base substitutions across the mU7 or hU7 hairpins; and single base substitutions across the natural U7 Sm or U1 Sm binding domains with a constant mU7 hairpin (Fig. 2a, Suppl. Table 2). A mutated Sm domain (SmMUT) with mU7 hairpin served as a negative control^9^. This SmOPT U7 library was cloned onto three gRNAs possessing a variety of structural features: 100.50 antisense against the RAB7A 3’UTR with a double hnRNP A1 domain, 100.50 antisense against the SNCA 3’UTR with no 5’ domain, and a 115.80 antisense against the GAPDH coding sequence possessing symmetrical loops at the −5 and +30 positions, previously shown to improve editing specificity^1^. All gRNAs were expressed using the mU7 promoter.

For the split-pool screen, plasmid libraries were first sequenced to match the gRNA with a 14N random “GuideID” barcode located in the GFP 5’UTR; across the six pools, 14870 total unique GuideIDs were identified (Suppl. Table 3). After these plasmid libraries were integrated at a single copy into Landing Pad 293T cells, 10 million cells were fixed and permeabilized. Custom primers annealed to and reverse transcribed GFP, RAB7A, SNCA, and GAPDH mRNA within the fixed cell. Each RT primer also included an additional 10N umi and an anchor sequence, which was then annealed to a Splint scaffold. Fixed, reverse-transcribed cells were then subjected to three rounds of Split-pool barcoding, wherein each cell may travel down any one of 216,000 potential combinations of wells, yet every transcript within that cell will be tagged identically. As a final round, 10,000 cells per well were dispensed across a PCR plate, lysed, and the released barcoded cDNA was amplified with dual index primers for Illumina sequencing. For each gRNA, transcripts from all cells possessing the corresponding 14N “GuideID” barcodes were aggregated, and the percent of RNA editing was calculated for RAB7A, SNCA, and GAPDH.

Our split-pool screen successfully detected RNA editing across all three endogenous transcripts (Fig. 2b). Furthermore, cell barcodes associated with editing of RAB7A, SNCA and GAPDH were exclusively linked to gRNAs designed to the corresponding target gene. A strong correlation was observed between the two replicate pools. Finally, negative control gRNAs possessing the non-functional SmMUT sequence did not show any editing. We concluded that the pooled screening method accurately reports RNA editing enabled by gRNAs, and proceeded to analyze the effects of SmOPT and hairpin mutations on editing.

### Split-pool screen identified SmOPT-U7 hairpin variants that improve RNA editing

Among all three transcripts, any mutation within the first seven nucleotides of the SmOPT sequence eliminated RNA editing, using either the mU7 or hU7 hairpin (Fig. 2c and Suppl. Fig. 3). Likewise, neither the wild type U1 Sm or U7 Sm binding domains showed RNA editing (consistent with the individual transfections in Suppl. Fig. 2), nor any of their mutated variants, except one which turns the U1 Sm directly into the SmOPT sequence (Suppl. Fig. 3). While less pronounced, some mutations along the U7 hairpin sequences also consistently reduced RNA editing on all targets (e.g. changing the first nucleotide from C/U to G, or changing the third nucleotide pair from G-C to A-U). Overall, the hU7 produced less RNA editing compared to the mU7 hairpin, despite HEK 293T being a human cell line.

**Fig. 3.**
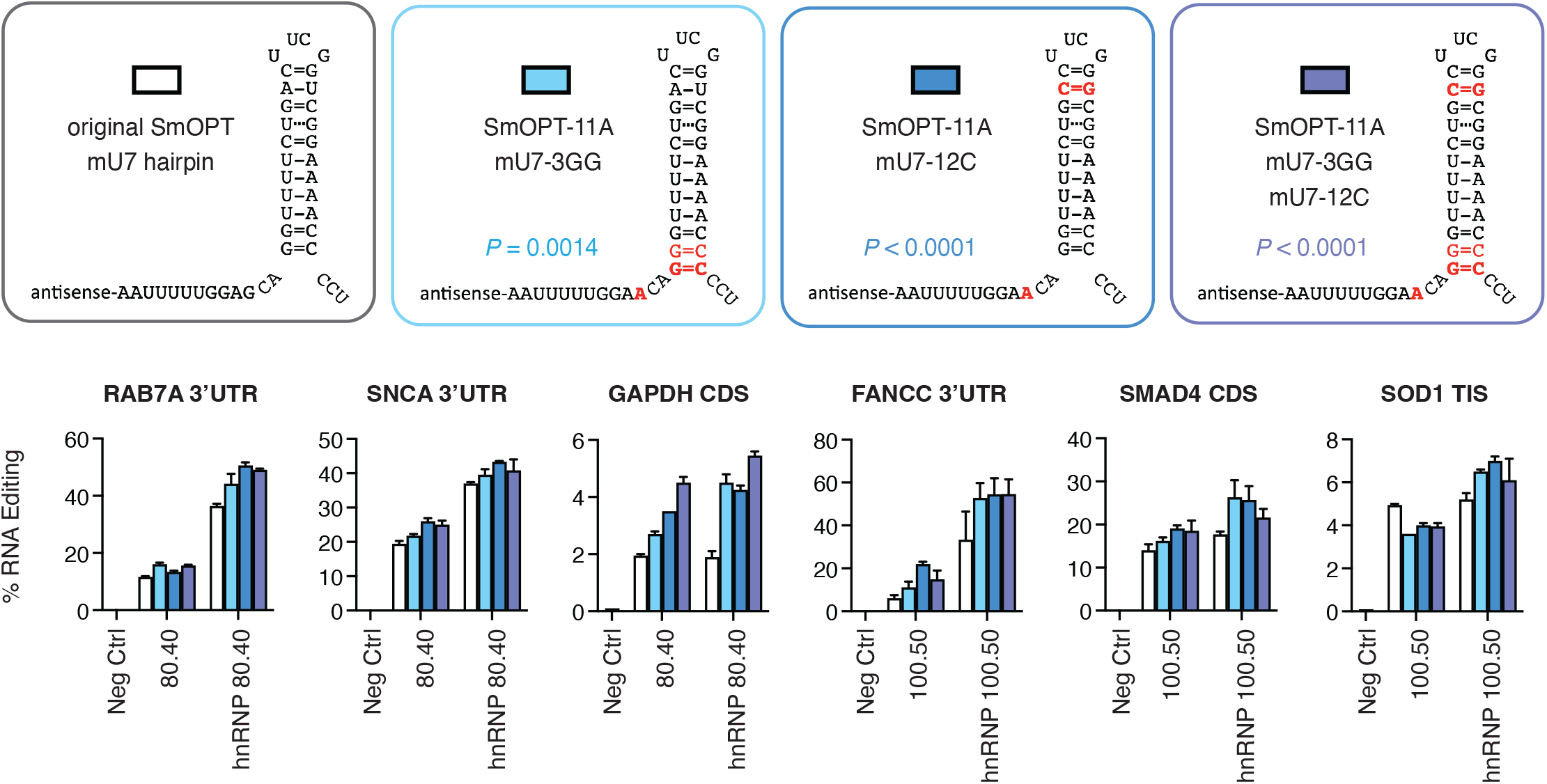
Novel SmOPT U7 variants can improve editing across multiple endogenous targets. Three combination variants along the SmOPT mU7 hairpin sequence (SmOPT-11A + mU7-3GG, SmOPT-11A + mU7-12C, and SmOPT-11A + mU7-3GG + mU7-12C) were cloned onto six antisense guide RNAs, with or without a 5’ double hnRNP A1 binding motif, and individually transfected into Landing Pad 293T cells. Following puromycin selection for successfully integrated single-copy guide RNA constructs, RNA was measured 13 days post transfection. Results from all 12 groups (6 gene targets +/-hnRNP) were analyzed in aggregate by two-way ANOVA; P values comparing each combination variant to the original SmOPT mU7 hairpin were calculated using Dunnett’s multiple comparisons test. Negative controls consist of samples where the transfected plasmid targeted a different gene.

Some mutations in the SmOPT or U7 hairpin sequence were found to increase RNA editing on all three transcripts. The top 23 mutations were individually cloned as plasmids and single-copy integrated into Landing Pad 293T cells (Fig. 2c). The individual validations largely confirmed the pooled screen results, with most variants outperforming the original SmOPT with mU7 hairpin. Although many mutations which inserted extra nucleotides before the SmOPT sequence increased editing, and were individually reproducible, these effects depended on the adjoining antisense sequence rather than a universal improvement to the SmOPT.

The top five individually validated mutations were then tested in every possible combination (2^5 = 32) on the three target genes (Suppl. Fig. 4). While many combinations increased editing compared to the original SmOPT mU7 hairpin sequence, three combination variants stood out: SmOPT-11A + mU7-3GG; SmOPT-11A + mU7-12C; and SmOPT-11A + mU7-3GG + mU7-12C. These combination variants were examined, via single-copy integration, on an additional set of antisense gRNAs, with or without the 5’ double hnRNP A1 binding motif (Fig. 3 and Suppl. Fig. 5). The FANCC 3’UTR, SMAD4 CDS, and SOD1 TIS were targeted with the same 100.50 antisense sequence in Figure 1.To confirm robustness of the improved SmOPT U7 hairpin sequences, the RAB7A 3’UTR, SNCA 3’UTR, and GAPDH CDS were now targeted with a shorter 80.40 antisense gRNA. Guide RNAs were expressed using the human U1 snRNA promoter^37^. Overall, the new SmOPT U7 hairpin variants significantly increased gRNA editing. A similar pattern was observed for antisense gRNAs targeting splice acceptor adenosines for DMD exon 71 or 74 skipping (Suppl. Fig. 6).

**Fig. 4.**
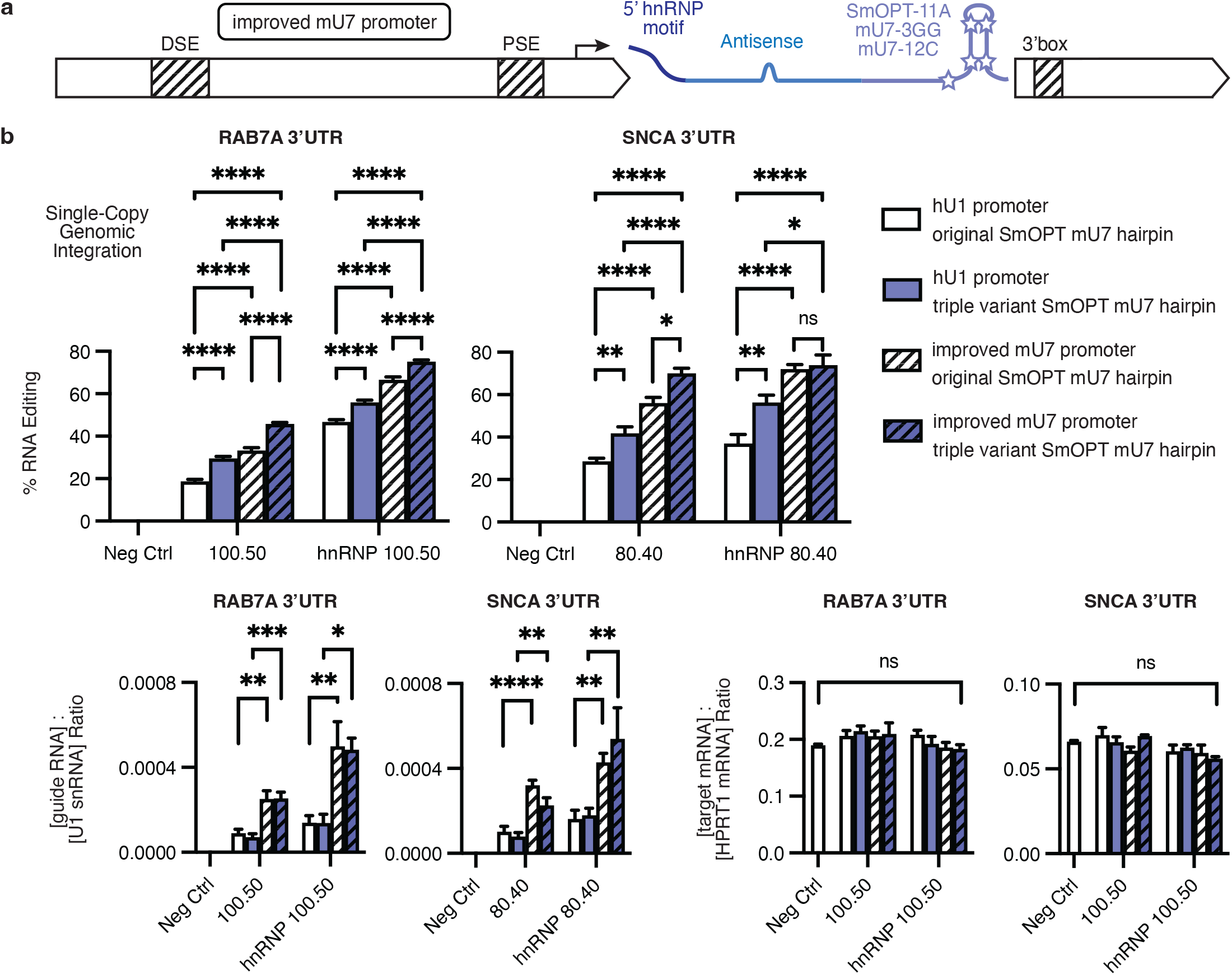
An improved synthetic U7 promoter further increases RNA editing and guide RNA expression. **a**, Arrangement of the synthetic guide RNA expression cassette; locations of the modified DSE, PSE, and 3’box elements within the mouse U7 promoter and terminator are highlighted. **b**, Antisense guide RNAs targeting the RAB7A 3’UTR (100.50) or SNCA 3’UTR (80.40), with or without a 5’ double hnRNP A1 binding motif, were cloned onto the original SmOPT U7 hairpin sequence or the triple variant (SmOPT-11A + mU7-3GG + mU7-12C). Guide RNAs were expressed using either the natural hU1 promoter and mU7 terminator or an improved synthetic mU7 cassette. Plasmids were individually transfected into Landing Pad 293T cells. RNA editing was measured 13 days post transfection, following puromycin selection for cells containing single-copy integrations of the guide RNA construct. Expression levels of guide RNA and target mRNA were measured using ddPCR and normalized to U1 snRNA or HPRT1 mRNA housekeeping genes, respectively (ratio of copies per µl). Results were analyzed in aggregate by three-way ANOVA: for RNA editing, P < 0.0001 for all three main effect variables (promoter, SmOPT U7 hairpin, and guide RNA); for guide RNA quantification, P < 0.0001 for the promoter and guide RNA main effect variables, but not significant for the SmOPT U7 hairpin. Asterisks indicate pairwise comparisons among each antisense guide RNA using one-way ANOVA with Tukey’s test.

**Fig. 5.**
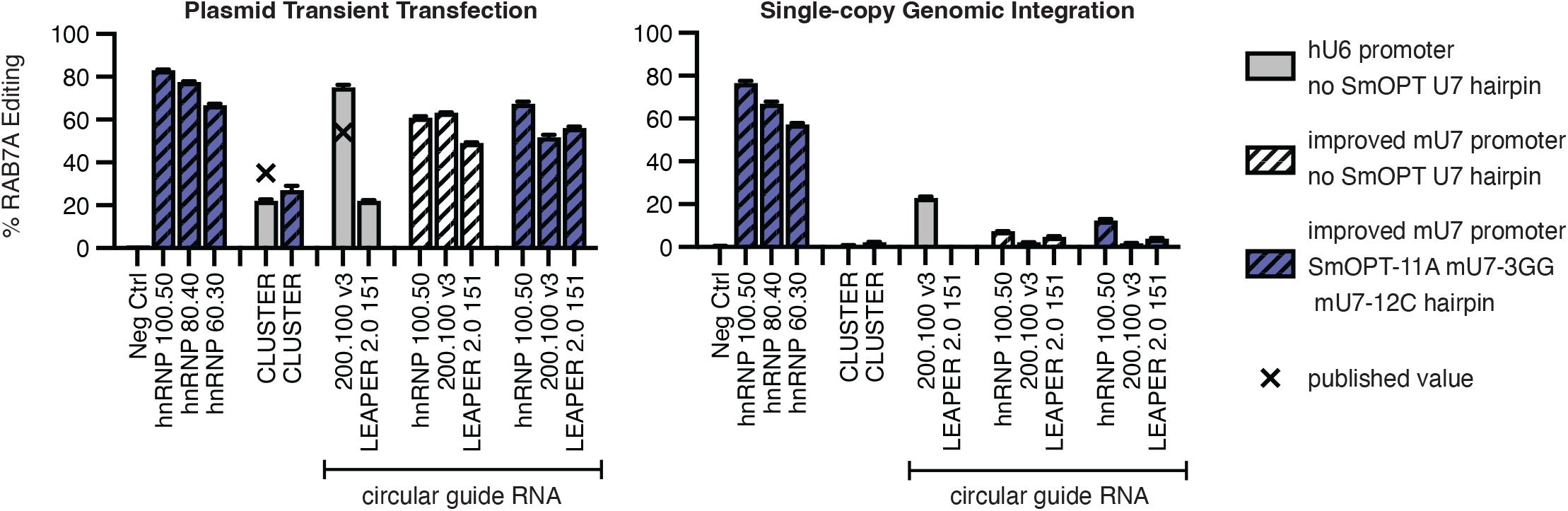
Comparison and combination with alternative gRNA design approaches. Antisense guide RNAs targeting the RAB7A 3’UTR (100.50, 80.40, or 60.30) with a 5’ double hnRNP A1 binding motif were compared to three previously published guide RNAs: CLUSTER 3xRS 19-13-11-20p8; cadRNA 200.100.loops.interspersed.v3; and LEAPER circ-arRNA151. Published RAB7A RNA editing values for transient transfection of 293 cells are marked. Guide RNAs were expressed using either the hU6 promoter (gray) or an improved synthetic mU7 promoter (striped). In some cases, the triple-variant SmOPT U7 hairpin was included on the antisense guide RNA (purple shaded). In other cases, the guide RNA sequences were flanked by circularizing ribozymes to form circular guide RNAs (as noted). Plasmids were individually transfected into Landing Pad 293T cells. For transient transfection, RNA editing was measured 2 days later. For single-copy genomic integration, RNA editing was measured 13 days post transfection, following puromycin selection for cells containing single-copy integrations of the guide RNA construct.

**Fig. 6.**
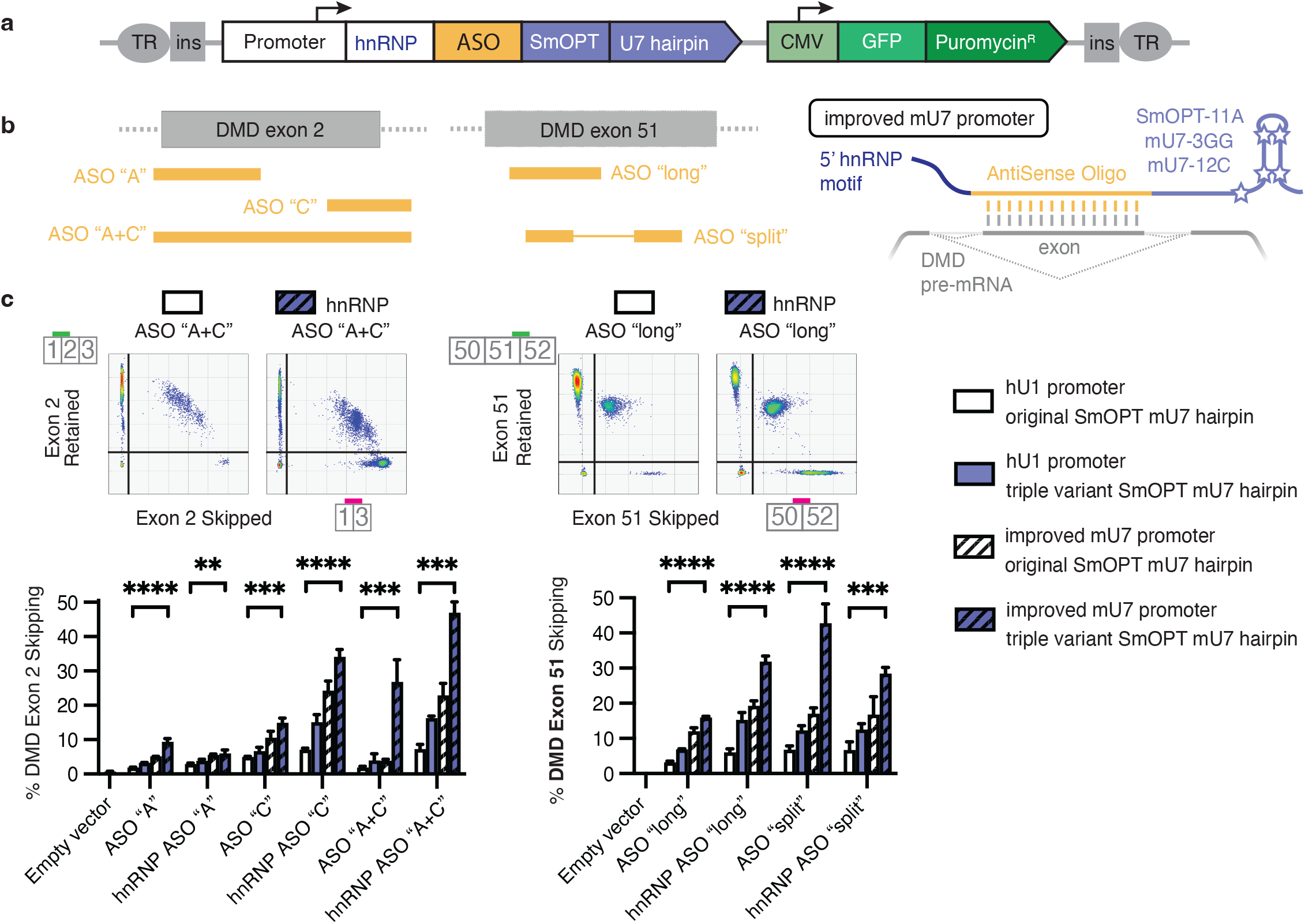
New SmOPT U7 hairpin variants also improve published ASOs for DMD exon skipping. **a**, Antisense Oligo RNA expression constructs alongside a constitutive GFP and puromycin resistance marker were flanked by insulator sequences (ins) and terminal repeats (TR) for the piggybac transposase. **b**, Placement of the ASO binding sites on DMD exon 2 or 51, which were expressed with or without a 5’ double hnRNP A1 binding motif. **c**, ASOs were expressed with the original SmOPT U7 hairpin (white) or the triple combination variant (SmOPT-11A + mU7-3GG + mU7-12C) (purple shaded), using either the natural hU1 promoter (open bars) and mU7 terminator or an improved synthetic mU7 cassette (striped bars). Constructs were randomly integrated into the genome of RD rhabdomyosarcoma cells using piggybac transposase, followed by 7 days of puromycin selection for successful integrants. GFP+ myoblasts were then differentiated for 10 days to induce expression of the full-length DMD Dp427m transcript. Exon skipping was measured using droplet digital PCR. Top: Representative ddPCR plots with probes specific for exon retained or exon skipped mRNA isoforms. Bottom: Percent exon skipping. Empty vector is a piggybac integration construct which contains the GFP and puromycin markers but lacks an antisense RNA. Results were analyzed in aggregate by three-way ANOVA with P < 0.0001 for all three main effect variables (promoter, SmOPT U7 hairpin, and guide RNA). Asterisks indicate pairwise comparisons among each antisense guide RNA using one-way ANOVA with Dunnett’s test.

### An improved U7 promoter further increases RNA editing

In addition to improving the SmOPT U7 hairpin structure, we reasoned that the gRNA promoter itself could be engineered for higher expression of gRNAs to increase editing performance from the same amount of delivered construct. While the pol II-type U7 or U1 promoters could mediate robust RNA editing, the pol III-type U6 promoter could not (Fig. 1). Even though RNA polymerase II also transcribes mRNA, the pol II snRNA genes lack a TATA box or polyadenylation sequence. Instead, full expression and maturation of pol II snRNAs requires Distal and Proximal Sequence Elements (DSE, PSE) within their promoters, and a 3’box signal before the termination sequence that recruits the Integrator complex for post-transcriptional processing^10,38^. We therefore tested synthetic versions of the mouse U7 snRNA promoter and terminator by duplicating or replacing the DSE, PSE, and 3’box with alternative motif sequences (Fig. 4a)^39^.

The top improved mU7 promoter increased RNA editing from a single integrated cassette when compared to the human U1 promoter, which had our top natural promoter. Upon a two-day plasmid transient transfection, everym combination of promoter and SmOPT mU7 hairpin reached a saturating level of RNA editing (Suppl. Fig. 7). However, when these constructs were tested at a single copy integration in Landing Pad 293T cells, the new synthetic mU7 promoter and the triple-variant SmOPT U7 hairpin (SmOPT-11A + mU7-3GG + mU7-12C) each independently significantly increased RNA editing (P < 0.0001 for each main effect variable) (Fig. 4b). Combined, they achieved 76% editing of the RAB7A 3’UTR and 68% editing of the SNCA 3’UTR. The improved promoter significantly increased gRNA expression, and target mRNA expression levels were unchanged.

**Fig. 7.**
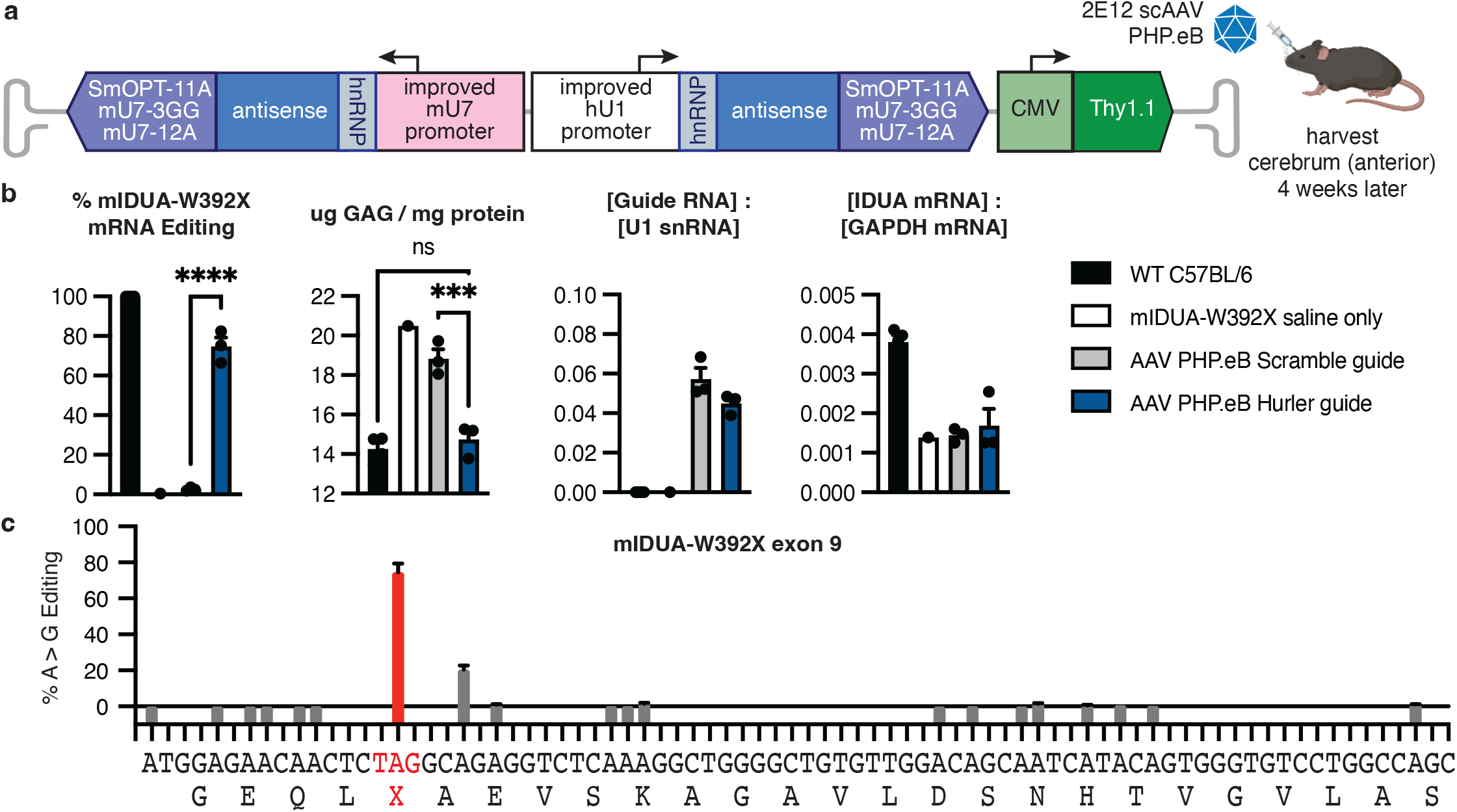
Improved SmOPT U7 gRNAs achieve high levels of RNA editing in Hurler syndrome mice. **a**, AAV vector used for guide RNA delivery. A 90.65 antisense Hurler guide RNA with –6 +30 loops targeting the mIDUA W392X pre-mRNA or a 100 nt antisense Scramble negative control were combined with a 5’ hnRNP domain and the triple variant SmOPT U7 hairpin. 2E12 scAAV PHP.eB were retroorbitally injected into mIDUA-W392X mutant mice. In addition, mIDUA-W392X mice injected with saline only or wild type C57BL/6 mice were included as controls. Brain tissue (anterior cerebrum) was harvested 4 weeks later. **b**, *In vivo* cerebral cortex samples were analyzed for: RNA editing of the target IDUA-W392X mutation; Glycosaminoglycan accumulation; Guide RNA expression level relative to the native U1 snRNA; and mIDUA transcript levels relative to GAPDH mRNA. Results were analyzed by one-way ANOVA with multiple pairwise comparisons. **c**, Bystander A > G editing along the region of mIDUA exon 9 targeted by the AAV PHP.eB Hurler guide. The W392X mutation is highlighted in red. Editing of the subsequent adenosine results in a synonymous GCA > GCG mutation (both alanine).

### The modified SmOPT U7 hairpin scaffold outperforms previous approaches to gene-encoded ADAR gRNAs

Several recent publications have described alternative approaches to ADAR gRNA design using gene-encoded vectors (no chemical modification) and endogenous ADAR enzyme levels (no risk of ADAR overexpression). We sought to compare – and possibly combine – our improved promoter and scaffold against these other methods. The CLUSTER approach appends a multivalent series of short antisense binding sequences following a 20.8 target sequence attached to an ADAR recruiting domain^40^. Two groups have published circular ADAR-recruiting gRNAs formed by flanking Twister ribozyme sequences, termed cadRNA and LEAPER 2.0^1,2^. These circular gRNAs contain a longer antisense domain (100-200 nt) interspersed with uridine deletions, G mismatches, or loops to reduce bystander editing. Fortunately, all of these publications included the RAB7A 3’UTR example target, tested by plasmid transient transfection of 293 cells, which allows for a direct comparison.

The top ADAR gRNAs against the RAB7A 3’UTR target were chosen from each publication: CLUSTER 3xRS 19-13-11-20p8 (35% editing); cadRNA 200.100.loops.interspersed.v3 (54% editing); and LEAPER circ-arRNA151 (editing value not listed). We expressed these gRNAs using either the hU6 promoter, as originally reported, or our improved mU7 promoter (Fig. 5). To investigate whether our scaffold could synergize with these other methods, we appended our triple-variant SmOPT U7 hairpin after the antisense gRNA (but within the circularizing ribozymes for circular gRNAs). For plasmid transient transfection, all constructs successfully edited the RAB7A 3’UTR, reproducing the published editing rates. Both CLUSTER and circular RNA constructs retained activity when expressed by the improved U7 promoter. The LEAPER circ-arRNA151 gained a particular boost from the modified mU7 promoter. This antisense sequence possesses a stretch of 5 uridines, which triggers premature termination from the U6 promoter; expressing this gRNA from the improved mU7 promoter raised RAB7A editing rates equivalent to the other circular RNAs. Conversely, transforming the hnRNP 100.50 antisense sequence into a circular gRNA did not increase RAB7A editing, with or without the triple-variant SmOPT U7 hairpin.

When these same constructs were tested under the stringent single-copy genomic integration condition, the linear hnRNP SmOPT U7 hairpin constructs retained most of their RNA editing activity. This property was mostly preserved when we reduced the length of the antisense footprint from 100nt down to 60nt, demonstrating a compatibility for shorter antisense sequences that were previously not possible with gene-encoded gRNAs and endogenous ADAR^1^. In contrast, the CLUSTER and circular gRNAs showed a dramatic decline in RNA editing under single copy levels, showing that these approaches require much higher levels of DNA construct than the modified SmOPT U7 approach to function efficiently.

### The novel promoter and SmOPT U7 hairpin also improve ASOs for exon skipping

We next sought to test whether our improved mU7 promoter, 5’ hnRNP domain, and triple-variant SmOPT U7 hairpin would further enhance published antisense oligos for exon skipping. Separate from ADAR RNA editing, multiple groups have attached the original SmOPT U7 hairpin onto short antisense RNA sequences designed to cover splicing elements and block the splicing machinery, resulting in that exon being excluded from the mature mRNA. In the case of Duchenne’s Muscular Dystrophy, an exon skipping strategy can restore translation of the DMD gene by removing duplicated or mutated exons, or shifting a codon reading frame. For DMD exon 2, we tested two published antisense sequences termed “A” and “C” which cover the splice acceptor and splice donor sites, respectively^22^. The scAAV9.U7-ACCA vector, which expresses two copies each of the “A” and “C” short RNAs, has shown efficacy and safety in mice, non-human primates, and human clinical trials (NCT04240314) up to 18 months post treatment^13,20,21^. In addition, we tested a 75 nt combined “A+C” antisense sequence that covers the entirety of DMD exon 2 (Fig. 6). For DMD exon 51, we tested two published antisense sequences, a 45 nt “long” and a discontinuous 43 nt “split”, which anneal within the exon and have also shown long-term efficacy in mice^18^.

We used piggybac transposase to randomly integrate the ASO SmOPT U7 constructs into the genome of RD human rhabdomyosarcoma cells along with a constitutively expressed GFP and puromycin resistance marker (Fig. 6a). After selection for successfully integrated cells, GFP^+^ myoblasts were differentiated for 10 days to induce expression of the full-length DMD transcript^41^. Differentiated myocytes exhibited even GFP expression levels, indicating equal levels of piggybac transduction (Suppl. Fig. 8). Exon skipping was measured by droplet digital PCR using probes specific for the exon-skipped or exon-retained isoforms (Fig. 6c).

Across all ten ASO variations, the improved synthetic mU7 promoter and triple-variant SmOPT U7 hairpin greatly increased the frequency of exon skipping over the original scaffold (P < 0.0001 for either variable), enabling up to 47% skipping for DMD exon 2 and 43% for DMD exon 51 (Fig. 6c). Adding the hnRNP domain also increased exon skipping for most – although not every – ASO. Combining all three improvements resulted in a 25-fold increase for exon 2 skipping (ASO “A+C”) and a 10-fold increase for exon 51 skipping (ASO “long”).

### Novel promoter and SmOPT U7 hairpin produce high RNA editing in the mouse brain

Finally, to demonstrate the utility of this platform *in vivo* for RNA editing, we chose the mouse IDUA-W392X model of Hurler syndrome, also used in previous ADAR studies^1,2^. This mouse model replicates a W402X premature stop codon mutation in human alpha-L-iduronidase, an enzyme essential for breaking down glycosaminoglycans. Hurler syndrome is currently treated with a weekly enzyme infusion to reduce GAG accumulation; however, this has limited effect in the central nervous system and does not cure the disease. Recent efforts have focused on engineering an IDUA fusion protein capable of crossing the blood-brain barrier and preventing cognitive decline^42^. In general, RNA editing approaches may be particularly suitable for the central nervous system (CNS), where highly edited natural transcripts such as GRIA2 are found, and high levels of ADAR2 expression are observed in addition to ADAR1^43^.

We adapted our Landing Pad system for single-copy integration to include a segment of the mouse IDUA gene (Suppl. Fig. 9a). The W392X mutation lies near the edge of exon 9, and circular gRNAs designed against the mRNA or pre-mRNA have been shown to work equally^2^. Thus, we first tested 100.50 antisense gRNAs targeting the mRNA or pre-mRNA. Again, the combination of a 5’ hnRNP domain, the improved synthetic mU7 promoter, and the triple-variant SmOPT U7 hairpin produced the highest RNA editing rates (Suppl. Fig. 9b). A second round of gRNA optimization shifting the length and position of the antisense region and introducing mismatch loops further increased editing (Suppl. Fig. 9c). A 90.65 antisense Hurler gRNA with loops targeting the mIDUA W392X pre-mRNA was chosen for *in vivo* testing.

AAV vectors were cloned containing two copies of the Hurler gRNA or a Scramble negative control gRNA; both vectors included 5’ hnRNP domains, improved mU7 and hU1 synthetic promoters, and triple-variant SmOPT U7 hairpins (Fig. 7a). To target the brain, 2E12 scAAV PHP.eB^44^ were injected systemically into mIDUA-W392X mutant mice. Additional controls were mutant mice injected with saline only, or wild type C57BL/6 mice. Four weeks post injection, the anterior part of the brain (largely cerebral cortex) was harvested. Mice receiving the Hurler gRNA showed 75% average editing of the W392X stop codon, which reduced GAG accumulation down to wild type levels (Fig. 7b). No exon skipping was observed (Suppl. Fig. 10). Hurler or Scramble AAV constructs generated equivalent levels of gRNA expression with no significant alteration in IDUA mRNA expression (we confirmed previous reports that W392X mice have lower IDUA mRNA expression levels compared to wild type mice^45^). ADAR editing with this gRNA design was fairly specific, with editing observed only at the target adenosine and one other adenosine. Notably, A > G editing at this bystander edited codon results in a synonymous mutation which continues to code for alanine (Fig. 7c). We conclude that the modified SmOPT U7 hairpin framework retains its remarkable potential for enabling high efficiency RNA editing when moving from *in vitro* to *in vivo* systems, even when delivered to deep brain regions with systemic injection of AAV.

## Discussion

Custom RNA editing through delivery of ADAR-harnessing gRNAs is a growing field in gene therapy, in part because using existing endogenous ADAR machinery eliminates the risks from delivery of foreign enzymes required for other forms of nucleic acid editing. Multiple groups have reported *in vivo* editing with endogenous ADAR, along with some recently initiated clinical trials^46^. The most clinically advanced approaches all deliver gRNAs directly as heavily chemically modified oligonucleotides. However, chemically synthesized ASOs require artificial nucleobases and continual redosing, and face challenges with tissue penetrance and biochemical toxicity^47^. In contrast, an AAV-delivered gene-encoded antisense gRNA is naturally expressed from within the target cell, and AAV-delivered payloads can persist for decades with a single treatment. A rapidly expanding repertoire of novel AAV serotypes now allows a variety of organs to be specifically targeted using a non-invasive systemic injection, e.g. PHP.eB for mouse brain, and emerging primate-targeting options44,48–50.

One challenge for AAV gene therapies, however, is that current transduction levels may be limited to one or a few copies of episome per cell. Our breakthrough scaffold achieves, for the first time, high RNA editing from a single DNA copy of construct (Fig. 5), which represents the minimal possible cellular dose of a gene therapy. Importantly, the high potency of RNA editing afforded by this new scaffold was observed *in vivo* as well as *in vitro*, and represents a substantial advance over previous approaches. For example, optimized circular gRNAs delivered with the same PHP.eB serotype to a Hurler syndome mouse model reported up to 30% editing in the brain stem, but only 5-10% editing in the cortex and other areas^2,51^. In contrast, our SmOPT U7 gRNAs enabled 75% editing in anterior cerebral cortex (Fig. 7).

Many of the advances reported here were enabled by our single copy integration system, which screens gRNAs under more stringent physiological levels and with more uniform expression and kinetics than a plasmid transient transfection. It also enabled a pooled gRNA library screen in cells, since editing of the endogenous transcript can be traced back to the cell’s single gRNA. The library mutagenesis screen across SmOPT U7 hairpin sequences not only revealed stronger variants, but also which residues are responsible for the gain in editing. The Optimized Sm domain, which binds the standard seven-protein Sm ring and leads to accumulation in the nucleus^52^, is required for activity, especially the first 7 nucleotides (AAUUUUU) (Fig. 2). In contrast, the wild type U7 Sm domain, which recruits two alternative Lsm proteins to the Sm ring, does not produce RNA editing. Mutating residues along the U7 hairpin – particularly near the front of the hairpin – also reduced RNA editing activity, although the detriment was less pronounced. The mutations enhancing RNA editing activity (mU7-3GG, 10A, and 12C), also increased the annealing strength of the hairpin. Notably, the mouse U7 hairpin consistently outperformed the human U7 hairpin, even though the screen was performed in human cells, indicating some sequence-specific hairpin preferences.

We found that the SmOPT U7 scaffold’s advantages for ASOs for exon skipping also apply to antisense gRNAs for ADAR editing (a finding also recently reported in a preprint^53^), indicating that gRNAs gain qualitative benefits from being processed through the snRNA maturation machinery. The processing and functional pathway of natural snRNAs are well-described in the literature^9,11,36,38,52^, and critically involve assembly with the Sm protein ring leading to shuttling of snRNAs into nuclear areas of pre-mRNA splicing. Notably, ADAR editing and splicing frequently occur co-transcriptionally, which likely explains why SmOPT U7 antisense RNAs enhance custom RNA editing as well as exon skipping. Given this finding, how is it possible that our improved gRNA constructs can skip exons when desired (DMD exons 2, 51, 71, and 74), yet avoid exon skipping when unwanted and instead enable editing (RAB7A, SNCA, IDUA)? The placement of the gRNA relative to the splicing elements on the target mRNA is likely an important factor. Altered splicing may be more likely in short coding exons versus a long 3’UTR. The usefulness of adding a 5’ hnRNP A1 binding domain also depended on the length and position of the gRNA – while it often enhanced RNA editing, in some cases it reduced editing. These variables may need to be empirically optimized for each target transcript, assessing both RNA editing and splicing outcomes. While here we used the split-pool assay to understand and improve the SmOPT and U7 hairpin sequences for gRNA editing, future applications of the method could allow gRNAs to be screened for RNA splicing, since the primers can be designed to capture whatever happens to the endogenous target mRNA. Our assay can currently screen a thousand gRNA constructs in parallel, which we anticipate could be scaled up further.

Before moving to clinical application of this technology, one powerful component to add will be the use of antisense sequences more heavily engineered to maximize target editing and minimize bystander editing. In this study, we employed a simple method of antisense design that uses a single A-C mismatch at the target base, sometimes combined with two mismatched loops up- and downstream of the target base. These design principles consistently enable efficient target editing and demonstrate reduced bystander editing (Fig. 7, Suppl. Fig. 11). Indeed in the Hurler mouse, we observed no off-target edits that would affect the protein sequence beyond the corrected mutation itself. However, this antisense design strategy may be insufficient for many clinical applications, where greater specificity is desired, and where relevant adenosines may appear in sequence contexts disfavored by ADAR^54^. Encouragingly, significant recent progress has been made in the design, high-throughput screening and machine learning-guided generation of ADAR gRNA sequences, wherein the secondary structure of a gRNA : target complex can be customized to focus and maximize ADAR activity on almost any desired adenosine^55,56^. Indeed the combination of highly-engineered antisense gRNA sequences and the SmOPT U7 framework is likely to be critical, since even a perfect gRNA : target RNA structure must be expressed in sufficient quantity at the right location inside target cells to enable its activity. We propose that combining the modified SmOPT U7 scaffold presented here with ML-engineered gRNA antisense sequences will unlock widespread clinical application of gene-encoded therapeutic ADAR editing.

In summary, our new U7 hairpin scaffold dramatically increases the potency of antisense RNAs so that they can work at one vector copy per cell – the lowest gene therapy dose possible – for ADAR editing. This builds on decades of success using AAV-delivered antisense SmOPT U7 hairpin RNAs, which have proven to be safe, long-lasting, and not perturb the transcriptome^12,13,18,21^. Beyond editing, making simple changes to the promoter and hairpin sequences of existing exon-skipping ASOs can improve their efficacy over 10-fold (Fig. 6). Using this new expression system, we achieved an unprecedented level of central nervous system (CNS) RNA editing *in vivo* (Fig. 7). Unlike delivery of oligonucleotide gRNAs, AAV delivery of gene-encoded gRNAs should have very high durability in non-dividing cells like neurons, even offering the possibility of one-time cures for serious diseases. Applications exist beyond correcting single point mutations: custom ADAR editing can also target stop codons, start codons, splice sites, regulatory elements, and even synthetic biology logic circuits^57^.

## Methods

### Cell culture

HEK293T cells and RD rhabdomyosarcoma cells (ATCC) were cultured in DMEM with Glutamax and 10% FCS (Thermo) according to ATCC instructions. Plasmids (1 µg transient or 0.5 µg integration) were transfected using TransIT-293 or TransIT-LT1 reagent (Mirus). After integration, cells were selected for 7 days with puromycin (Sigma). RD cells were differentiated for 10 days on collagen-coated plates using DMEM supplemented with 2% horse serum, 0.1 µM TPA, and 5 µM GSK126 (Sigma).

To generate the Landing Pad cell line, first, an attP1 site was inserted into the AAVS1 genomic locus of HEK293T cells using a Cas9-expressing plasmid^58^. Single cell clones were screened by PCR for heterozygous knockins. Then, a plasmid containing the Landing Pad cassette and an attB1 site (Fig. 1c) was incorporated into these cells using a BxB1 integrase-expressing plasmid^30,31^. Cells containing the Landing Pad cassette were selected using blasticidin (Sigma) and monitored for FusionRed expression. For single copy integration experiments, cells were cultured with 2 µM doxycycline (Sigma) to induce expression of the BxB1 integrase.

### Molecular biology

Plasmids were purified by anion-exchange columns (Qiagen, Macherey-Nagel) and quantified on a Qubit fluorometer (Invitrogen). RNA was purified using the Qiagen RNeasy Mini kit with DNase (cell culture) or RNeasy Plus Universal Mini kit (organs). cDNA was reverse transcribed using oligo d(T)23VN, random hexamers, or custom primers using SuperScript IV (Thermo). Target sequences were PCR amplified using KAPA HiFi HotStart DNA polymerase, labeled with NEB Next dual index primers, and sequenced on Illumina iSeq or MiSeq platforms. Exon skipping and gene expression were quantified using the Bio-Rad QX200 ddPCR Supermix for Probes (no dUTP). Primer (IDT) and gRNA sequences are listed in Suppl. Table 1.

### Single cell sequencing

A library containing 252 mutations across the Sm binding and U7 hairpin sequences was synthesized (Twist Biosciences, Suppl. Table 2) and cloned onto three antisense gRNA plasmids in duplicate pools (NEB Golden Gate Assembly kit), each comprising approximately 3000 bacterial colonies. Plasmid libraries were sequenced to confirm diversity and match the gRNA with a 14N random barcode before the GFP gene (Fig. 2a, Suppl. Table 3.)

The Split pool barcoding method was adapted from O’Huallachain, et al.^35^ After formaldehyde fixation and methanol permeabilization, cDNA was reverse transcribed within the fixed cell using phosphorylated custom primers and SuperScript IV (Thermo). To retain cDNA within the cell, amino-allyl dUTP was included in the reverse transcription reaction, and the resulting cDNA was covalently attached to the cell with a DTSSP crosslinker (Thermo). Splint scaffolds were annealed to the RT primers, and then cells were subjected to three rounds of Split-pool barcode ligation (NEB T4 DNA ligase). Finally, aliquots containing 10,000 cells each were broken apart with proteinase K, RNAse H, and DTT (to reduce the cross-linker) (NEB, Thermo); and the released barcoded cDNA was amplified with dual index primers for Illumina sequencing. Our computational analysis pipeline first deduplicated sequencing reads by umi (filtering out any reads appearing less than thrice) and grouped transcripts according to their ligated series of subcodes “CellID”. To be included, “cells” needed at least 5 unique (by umi) transcripts overall, at least 3 unique GFP transcripts, and with at least 77% of the GFP transcripts sharing the same 14N “GuideID”. For each gRNA, transcripts from all “cells” possessing the corresponding “GuideIDs” were aggregated, and the percent of RNA editing was calculated for each gene target.

### AAV vectors

AAV PHP.eB was produced by triple transfection of plasmids encoding the recombinant transfer vector, pHelper, and PHP.eB RepCap^44^ into HEK293T cells using PEI (Polysciences). Virus was harvested, purified using an iodixanol density gradient ultracentrifugation method, concentrated using ultrafiltration spin columns (Cytiva), resuspended in buffered saline, and quantified using the Takara AAVpro Titration kit.

### Mouse experiments

Male C57BL/6J and IDUA-W392X (B6.129S-*Idua*^*tm1*.*1Kmke*^/J) mice^45^ were purchased from Jackson Labs. 5-6-week-old mice were injected retro-orbitally with 2.0 × 10^12^ AAV PHP.eB or a saline-only control; brain tissue was harvested 28 days later. All animal procedures were performed in accordance with protocol S16003 approved by the Institutional Animal Care and Use Committee of the University of California, San Diego.

### GAG assay

The GAG assay was performed as previously described^1^. Briefly, mouse tissues were homogenized in PBS with 0.05% Tween-20 using Qiagen QiaShredder columns. Samples were digested with 0.1 mg/ml proteinase K (NEB) at 56°C for 3 hours and then 95°C for 10 min to inactivate the enzyme. Soluble protein was clarified by centrifugation and quantified using the Pierce Quantitative Colorimetric Peptide Assay (Thermo). GAG was measured using the Blyscan GAG assay kit (Biocolor).

## Acknowledgements

We thank the Shape Therapeutics Vector Production Team for generating AAV; Nicole Enger for assisting the synthetic promoter development; and Dhruva Katrekar and Jacob Tome for technical advice.

## Author contributions

S.M. Byrne conceived the project with assistance from A.W.B. and P.M. S.M. Byrne devised gRNAs with the hnRNP and SmOPT U7 domains and adapted the split-pool library screening method. S.M. Burleigh designed and developed the synthetic promoter and AAV constructs. R.F. designed and built the Landing Pad system for single-copy integrated barcoded gRNAs. Y.J. developed the computational analysis pipeline. Y.S. advised on the placement of mismatch loops. S.M. Byrne and R.P. designed and performed experiments to test individual gRNAs. J.R. and A.P. conducted mouse experiments. S.M. Byrne, A.W.B., and P.M. wrote the paper with input from other authors.

## Grant acknowledgement

This work was generously supported by Institutional Funds.

---

**Supplementary Fig. 1.**
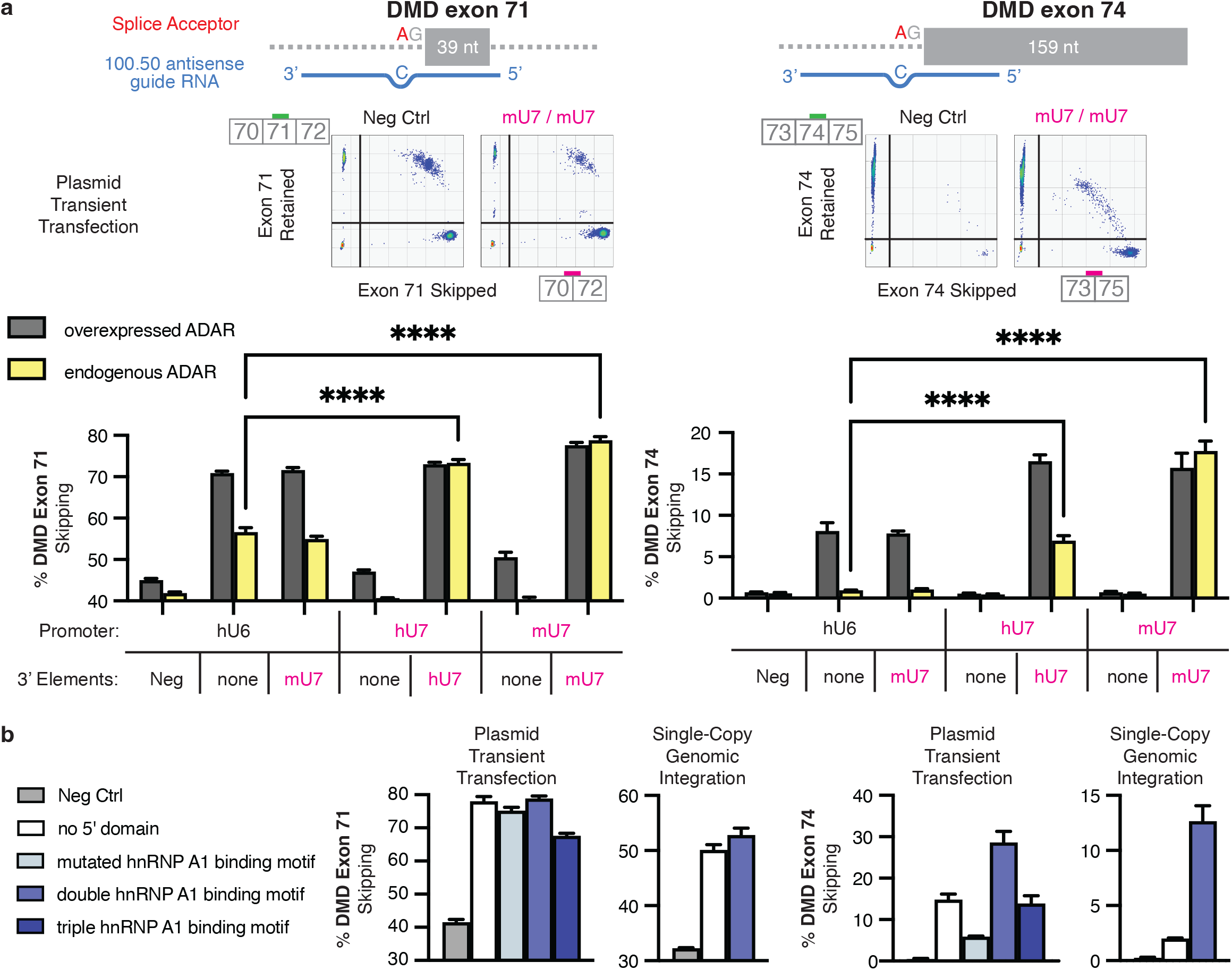
Antisense guide RNAs targeting splice acceptor adenosines for ADAR editing can mediate exon skipping. **a)** 100 nt antisense guide RNAs targeting adenosines in the splice acceptor site before DMD exon 71 or 74 were expressed using the hU6, hU7, or mU7 promoters, either alone or possessing an additional SmOPT sequence and human or mouse U7 hairpin on the 3’end. Plasmids also contained the CMV promoter to overexpress ADAR2 (gray bars) or GFP (endogenous ADAR, yellow bars). RNA was measured from HEK293T cells 2 days post transfection. The percent of DMD exon skipping was quantified using droplet digital PCR probes specific for either the exon-retained or exon-skipped transcript. HEK293T cells ordinarily express both the Dp71 and Dp71a (exon 71 skipped) DMD mRNA isoforms; thus, the baseline level of exon 71 skipping is around 30-50%. **b)** Sequences containing two or three copies of the hnRNP A1 binding motif were added onto the 5’ end of antisense guide RNAs expressed using a mouse U7 promoter and possessing a SmOPT mU7 hairpin. Plasmids were individually transfected into Landing Pad 293T cells. For transient transfection, RNA editing was measured 2 days later. For single-copy genomic integration, RNA editing was measured 13 days post transfection, following puromycin selection for cells containing a single copy of the guide RNA construct. Negative controls were measured from samples where the transfected plasmid targeted a different gene. Significance was calculated by Mixed-effects analysis using Dunnett’s multiple comparisons test.

**Supplementary Fig. 2.**
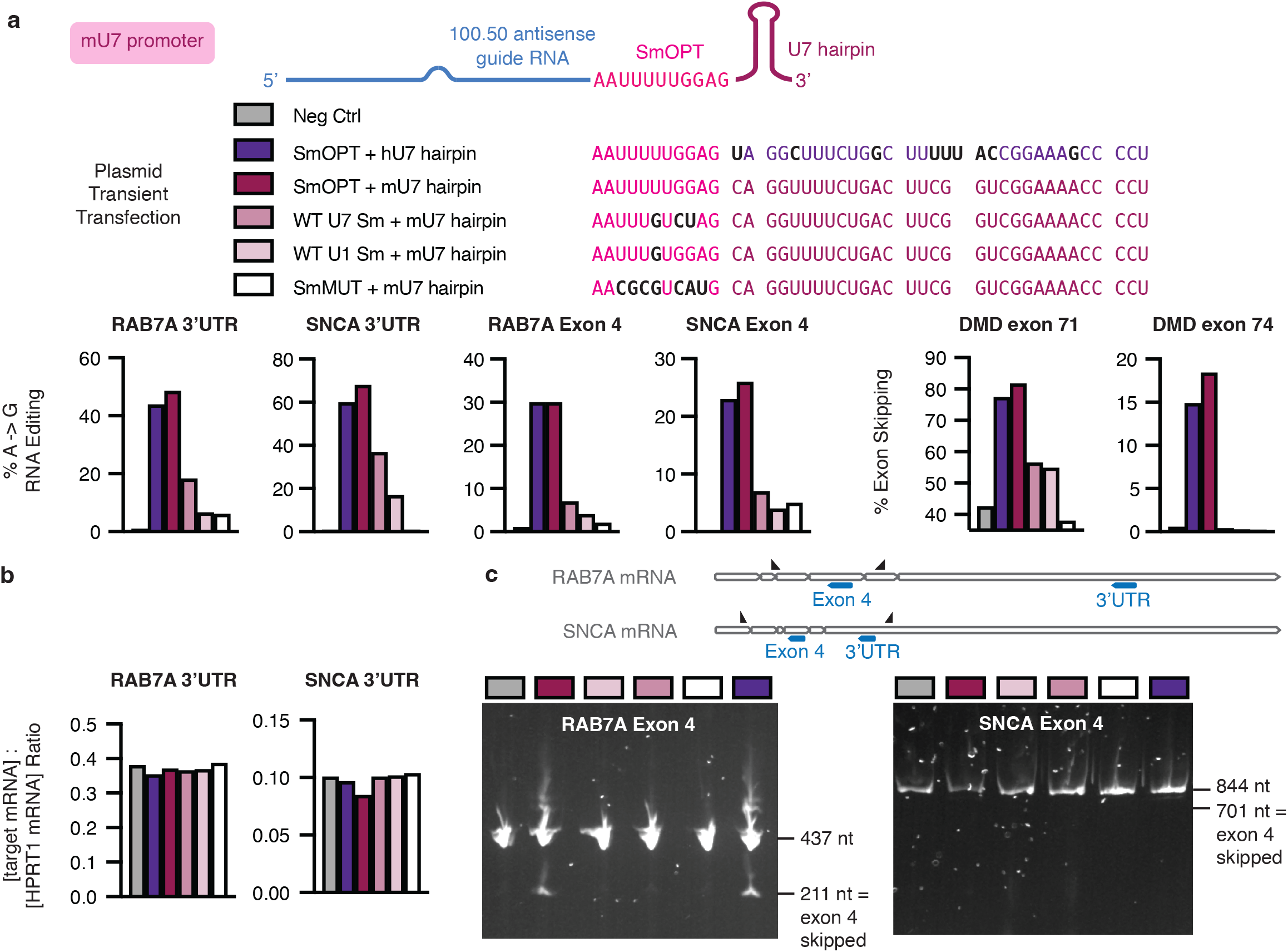
The Optimal Sm domain (SmOPT) is required for RNA editing activity. **a**, 100 nt antisense guide RNAs targeting adenosines in RAB7A 3’UTR or exon 4, SNCA 3’UTR or exon 4, or the splice acceptor site before DMD exon 71 or 74 were expressed using the mU7 promoter. Guide RNAs were attached to different Sm binding domains and either the human or mouse U7 hairpin. Plasmid constructs were transfected into HEK293T cells; RNA was purified 2 days later. The percent of editing or exon skipping for each target is shown. **b**, Expression levels of target mRNA were measured using ddPCR and normalized to HPRT1 mRNA (ratio of copies per µl). **c**, cDNA was PCR amplified using primers flanking RAB7A exon 4 or SNCA exon 4 (triangles) and run on an agarose gel.

**Supplementary Fig. 3.**
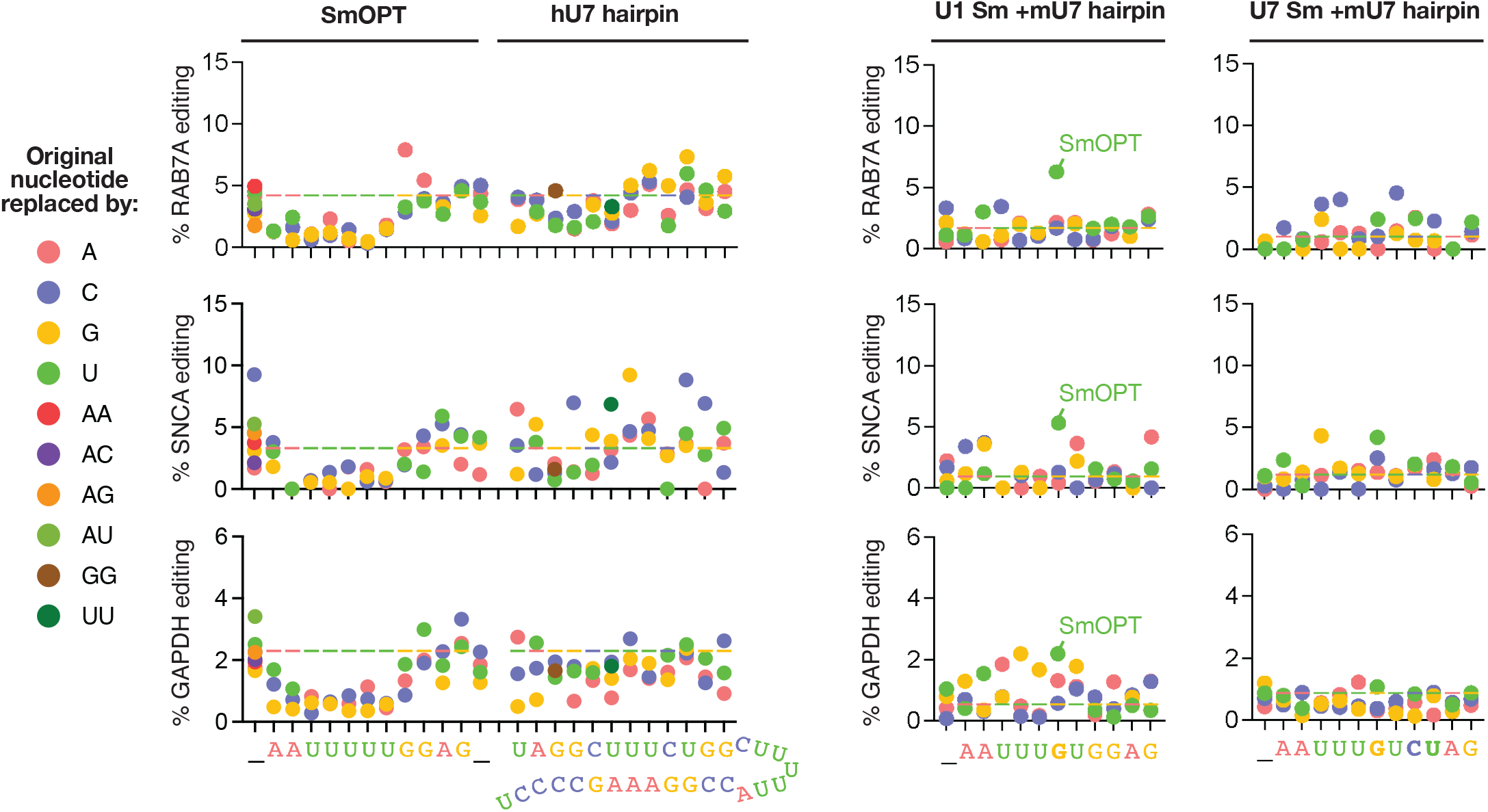
Additional results of the split-pool screen. For each gene target in the SmOPT U7 library screen, the effect on RNA editing of single nucleotide substitutions across the SmOPT sequence or paired substitutions across the hU7 hairpin is shown. Underscores indicate locations where extra nucleotides were inserted. The multicolor dashed line marks the percent editing from the original SmOPT hU7 hairpin construct. The effect on RNA editing of single nucleotide substitutions across the natural U1 or U7 Sm binding domains with an intact mU7 hairpin is also shown. Replacing the sixth nucleotide in the natural U1 Sm binding domain from G to U produces the original SmOPT sequence.

**Supplementary Fig. 4.**
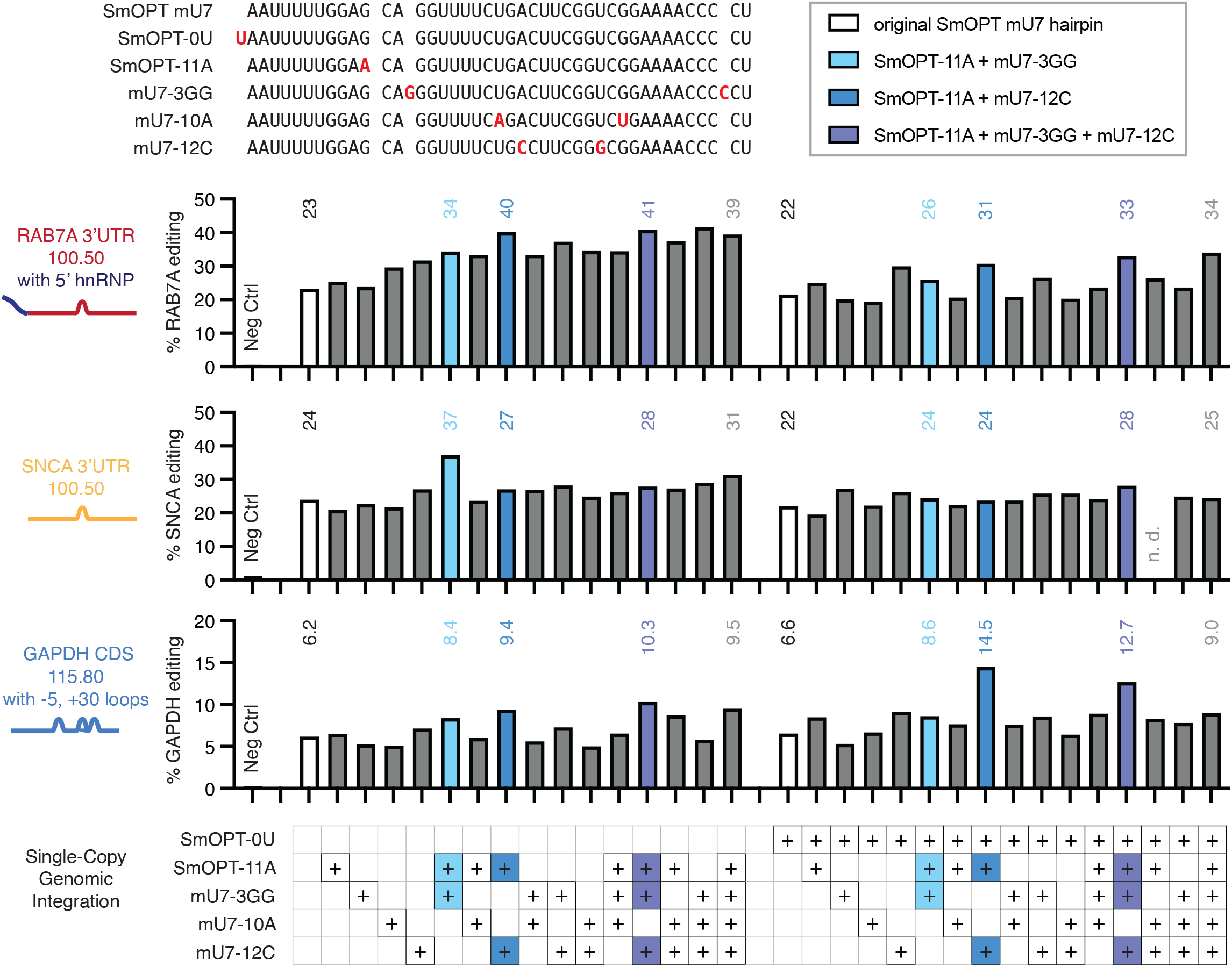
Combinations of top variants further boost editing. The top five individually-validated mutation variants along the SmOPT mU7 hairpin sequence (SmOPT-0U, SmOPT-11A, mU7-3GG, mU7-10A, mU7-12C, listed above), were cloned in every possible combination (2^5 = 32) onto three antisense guide RNAs and individually transfected into Landing Pad 293T cells. Following puromycin selection for successfully integrated single-copy guide RNA constructs, RNA was measured 13 days after the initial plasmid transfection. Selected combinations are highlighted with the percent of RNA editing noted above. n.d. = not determined. Negative controls were measured from samples where the transfected plasmid targeted a different gene.

**Supplementary Fig. 5.**
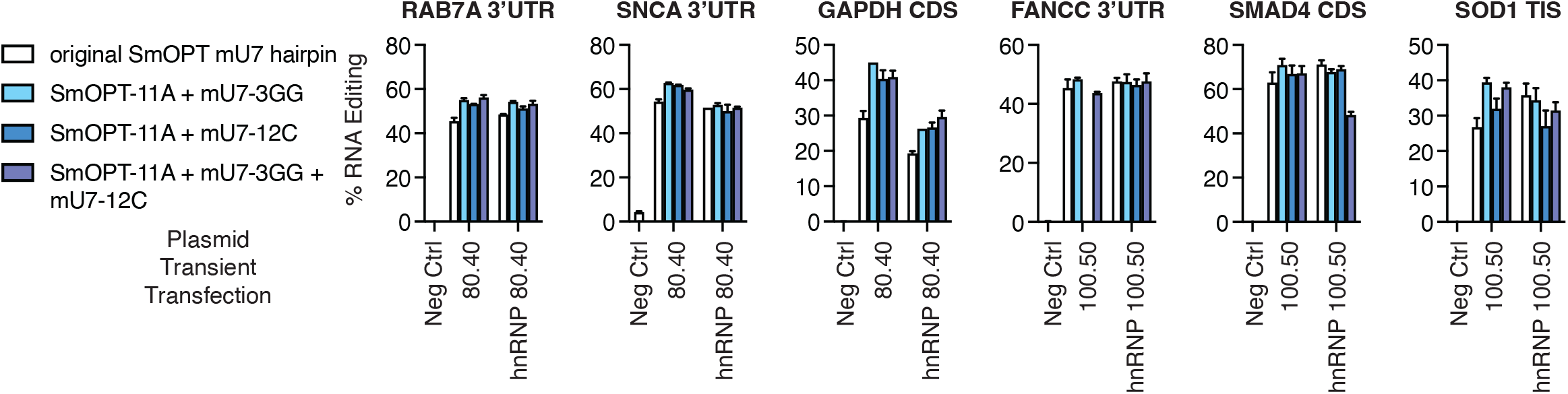
Transient transfection data of three new SmOPT U7 combination variants. Three combination variants along the SmOPT mU7 hairpin sequence (SmOPT-11A + mU7-3GG, SmOPT-11A + mU7-12C, and SmOPT-11A + mU7-3GG + mU7-12C) were cloned onto six antisense guide RNAs, with or without a 5’ double hnRNP A1 binding motif, and individually transfected into Landing Pad 293T cells. RNA editing was measured 2 days post transfection. (Not determined: FANCC 100.50 SmOPT-11A + mU7-12C). Negative controls consist of samples where the transfected plasmid targeted a different gene.

**Supplementary Fig. 6.**
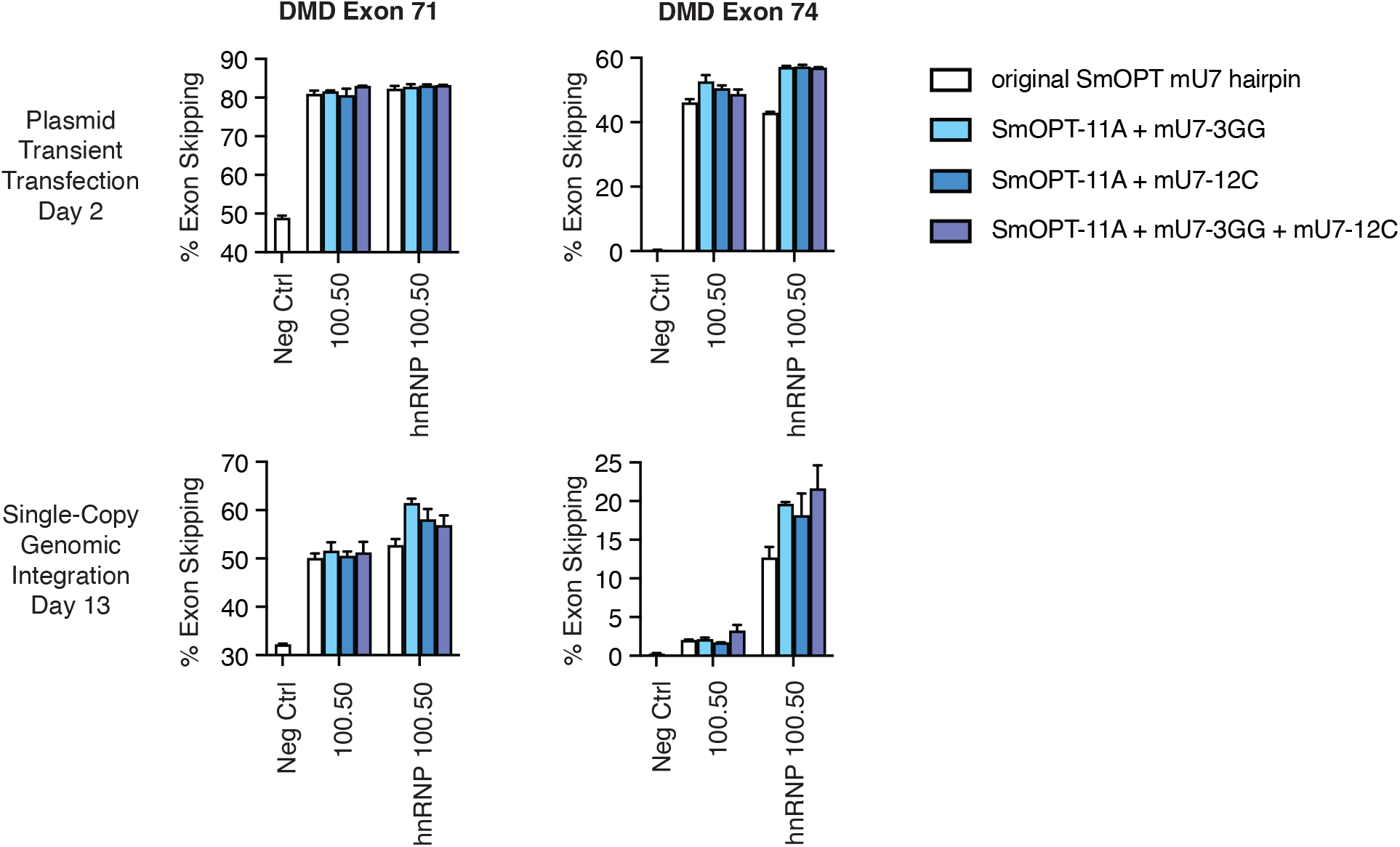
New SmOPT U7 combinations can also improve antisense gRNAs for DMD exon skipping. Three combination variants along the SmOPT mU7 hairpin sequence (SmOPT-11A + mU7-3GG, SmOPT-11A + mU7-12C, and SmOPT-11A + mU7-3GG + mU7-12C) were cloned onto two antisense guide RNAs targeting the splice acceptor adenosines of DMD exons 71 or 74, with or without a 5’ double hnRNP A1 binding motif, and individually transfected into Landing Pad 293T cells. For transient transfection, RNA was measured 2 days later. Following puromycin selection for successfully integrated single-copy guide RNA constructs, RNA was measured 13 days post transfection. The percent of DMD exon skipping was quantified using droplet digital PCR probes specific for either the exon-retained or exon-skipped transcript. HEK293T cells ordinarily express both the Dp71 and Dp71a (exon 71 skipped) DMD mRNA isoforms; thus, the baseline level of exon 71 skipping is around 30-50%. Negative controls consist of samples where the transfected plasmid targeted a different gene.

**Supplementary Fig. 7.**
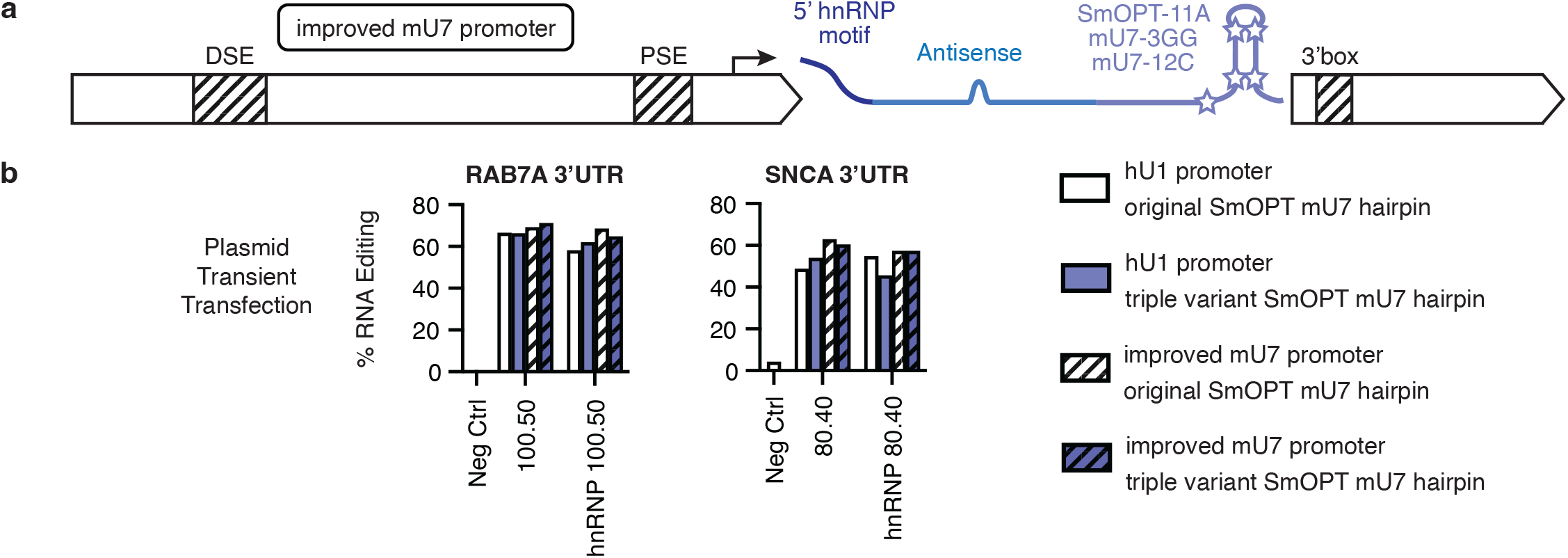
Transient transfection of new triple variant SmOPT U7 hairpin with improved U7 synthetic promoter. **a**, Arrangement of the synthetic guide RNA expression cassette; locations of the modified DSE, PSE, and 3’box elements within the mouse U7 promoter and terminator are highlighted. b, Antisense guide RNAs targeting the RAB7A 3’UTR (100.50) or SNCA 3’UTR (80.40), with or without a 5’ double hnRNP A1 binding motif, were cloned onto the original SmOPT U7 hairpin sequence or the triple variant (SmOPT-11A + mU7-3GG + mU7-12C). Guide RNAs were expressed using either the natural hU1 promoter and mU7 terminator or an improved synthetic mU7 cassette. Plasmids were individually transfected into Landing Pad 293T cells. RNA editing was measured 2 days later.

**Supplementary Fig. 8.**
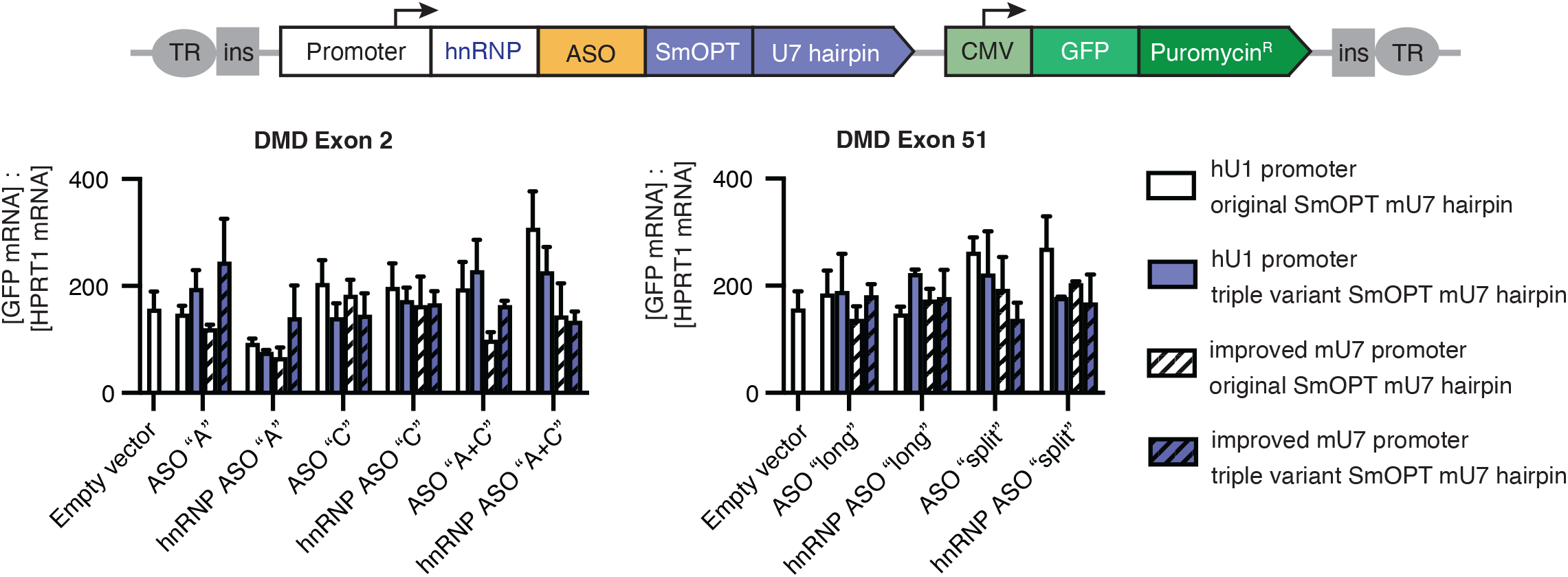
Even GFP expression across piggybac-integrated differentiated myoblasts. Constructs expressing Antisense Oligos targeting DMD exon 2 or 51, along with a constitutive GFP and puromycin resistance marker, were randomly integrated into the genome of RD rhabdomyosarcoma cells using piggybac transposase, as described in Figure 6. After 7 days of puromycin selection for successful integrants, GFP+ myoblasts were then differentiated for 10 days using TPA and GSK126 to induce expression of the full-length DMD Dp427m transcript. The ratio of GFP mRNA to HPRT1 mRNA was quantified using droplet digital PCR. Empty vector is a piggybac integration construct which contains the GFP and puromycin markers but lacks an antisense RNA.

**Supplementary Fig. 9.**
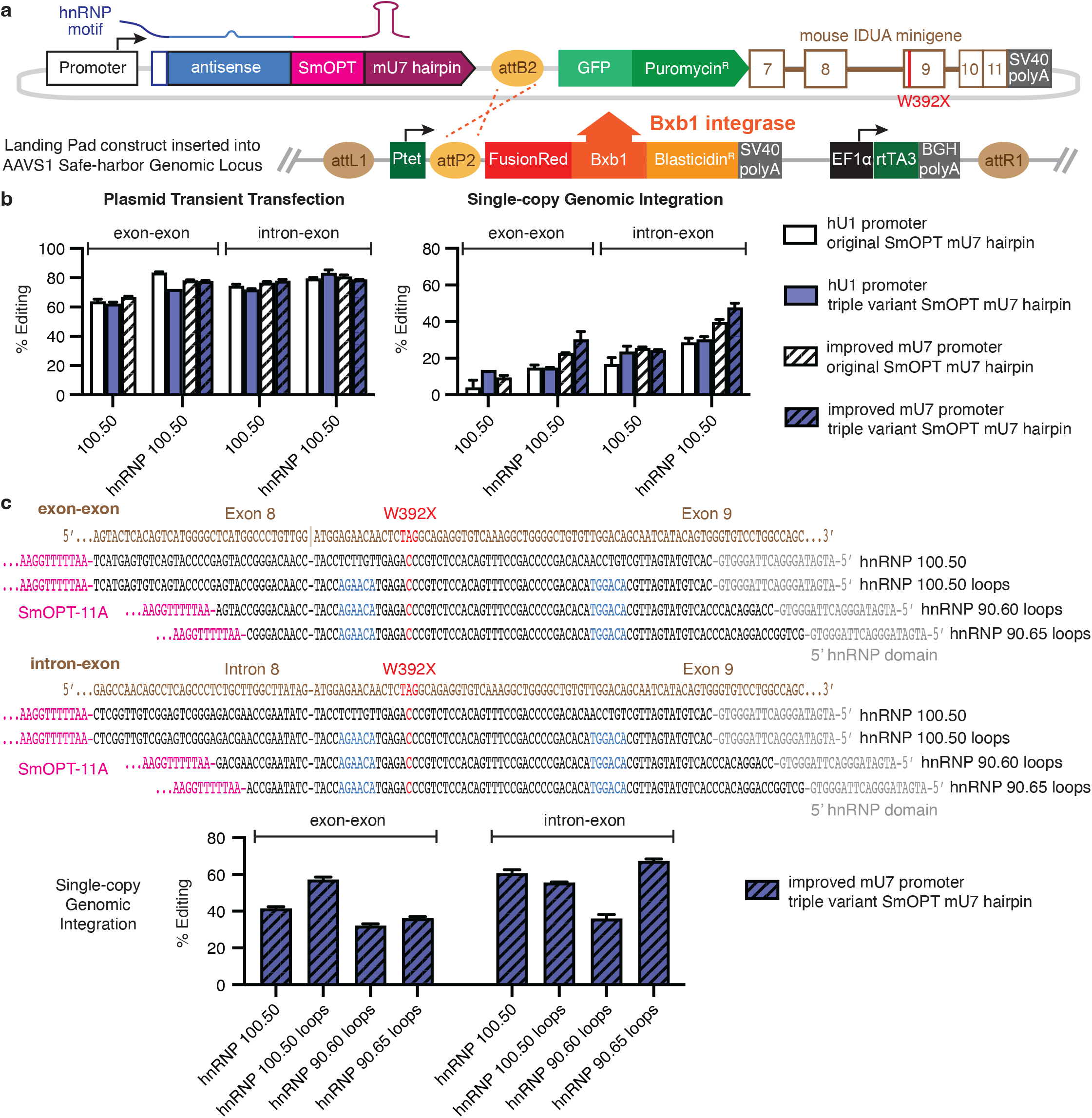
Optimization of Hurler guide RNAs. **a**, Construct for optimizing Hurler guide RNAs. A section of the mouse IDUA-W392X gene was inserted as a 3’UTR following the GFP and puromycin-resistance selection markers into the plasmid used for single-copy integration of guide RNAs described in Figure 1C. These plasmids were individually transfected into a Landing Pad HEK293T cell line possessing a doxycycline-inducible BxB1 integrase as previously described. For transient transfection, RNA editing of the mIDUA-W392X minigene was measured 2 days post transfection. For single-copy genomic integration, RNA editing was measured 13 days post transfection, following puromycin selection for cells containing the guide RNA construct. **b**, Antisense guide RNAs (100.50) targeting the mIDUA-W392X exon-exon (mRNA) or intron-exon (pre-mRNA), with or without a 5’ hnRNP A1 binding domain, were cloned onto the original SmOPT U7 hairpin sequence or the triple combination variant (SmOPT-11A + mU7-3GG + mU7-12C). Guide RNAs were expressed using either the natural hU1 promoter and mU7 terminator or an improved synthetic mU7 expression cassette. **c**, Variations of the exon-exon or intron-exon antisense guide RNAs were generated with mismatch loops and shifted annealing footprints. All guide RNAs contained the 5’ hnRNP A1 binding domain, the triple variant SmOPT mU7 hairpin, and the improved synthetic mU7 promoter. The intron-exon hnRNP 90.65 loops guide RNA was chosen for further *in vivo* testing.

**Supplementary Fig. 10.**
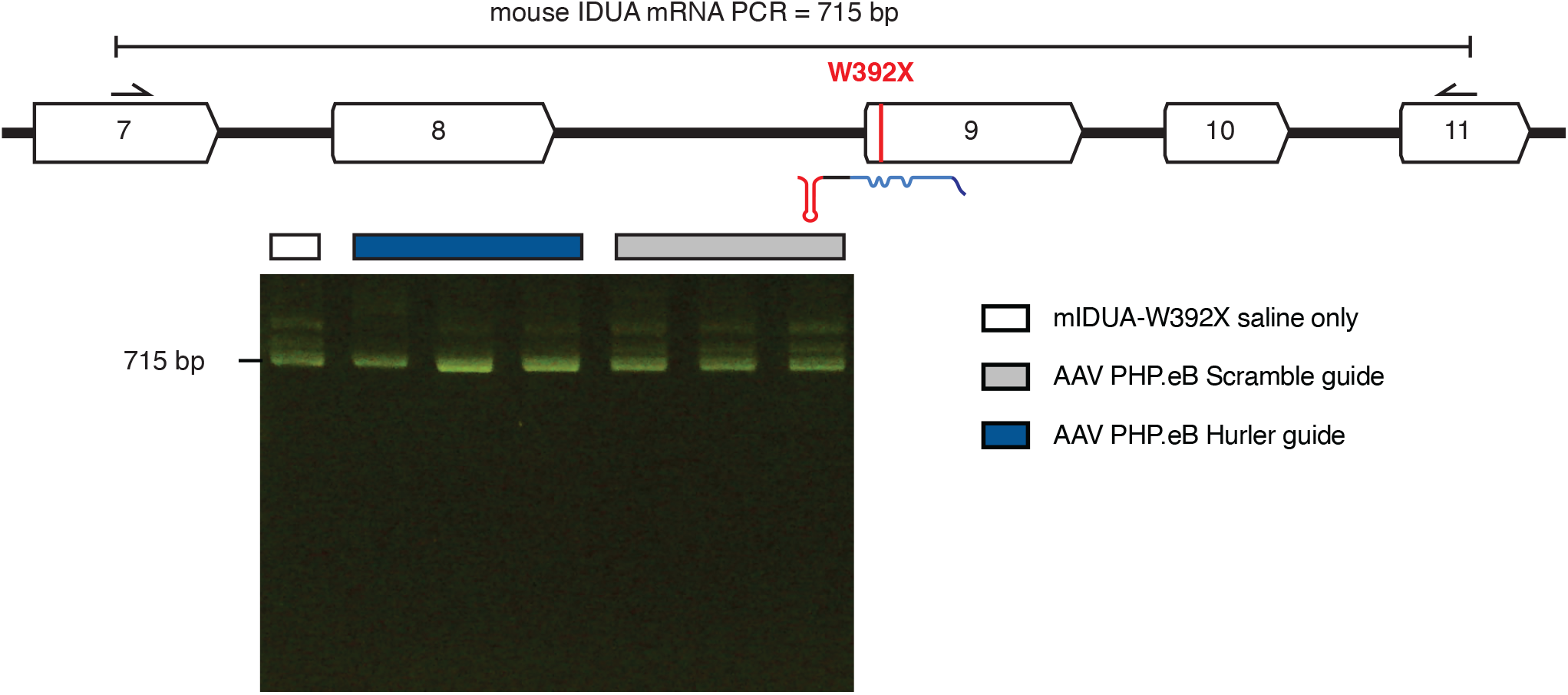
Improved SmOPT U7 guide RNAs do not cause exon skipping in Hurler syndrome mice. A 90.65 antisense Hurler guide RNA with –6 +30 loops targeting the mIDUA W392X pre-mRNA or a 100 nt antisense Scramble negative control were combined with a 5’ hnRNP domain and the triple variant SmOPT U7 hairpin. 2E12 scAAV PHP.eB or saline only was retroorbitally injected into mIDUA-W392X mutant mice. Brain tissue (anterior cerebrum) was harvested 4 weeks later. cDNA was PCR amplified using primers spanning mIDUA exons 7 and 11 and run on an agarose gel.

**Supplementary Fig. 11.**
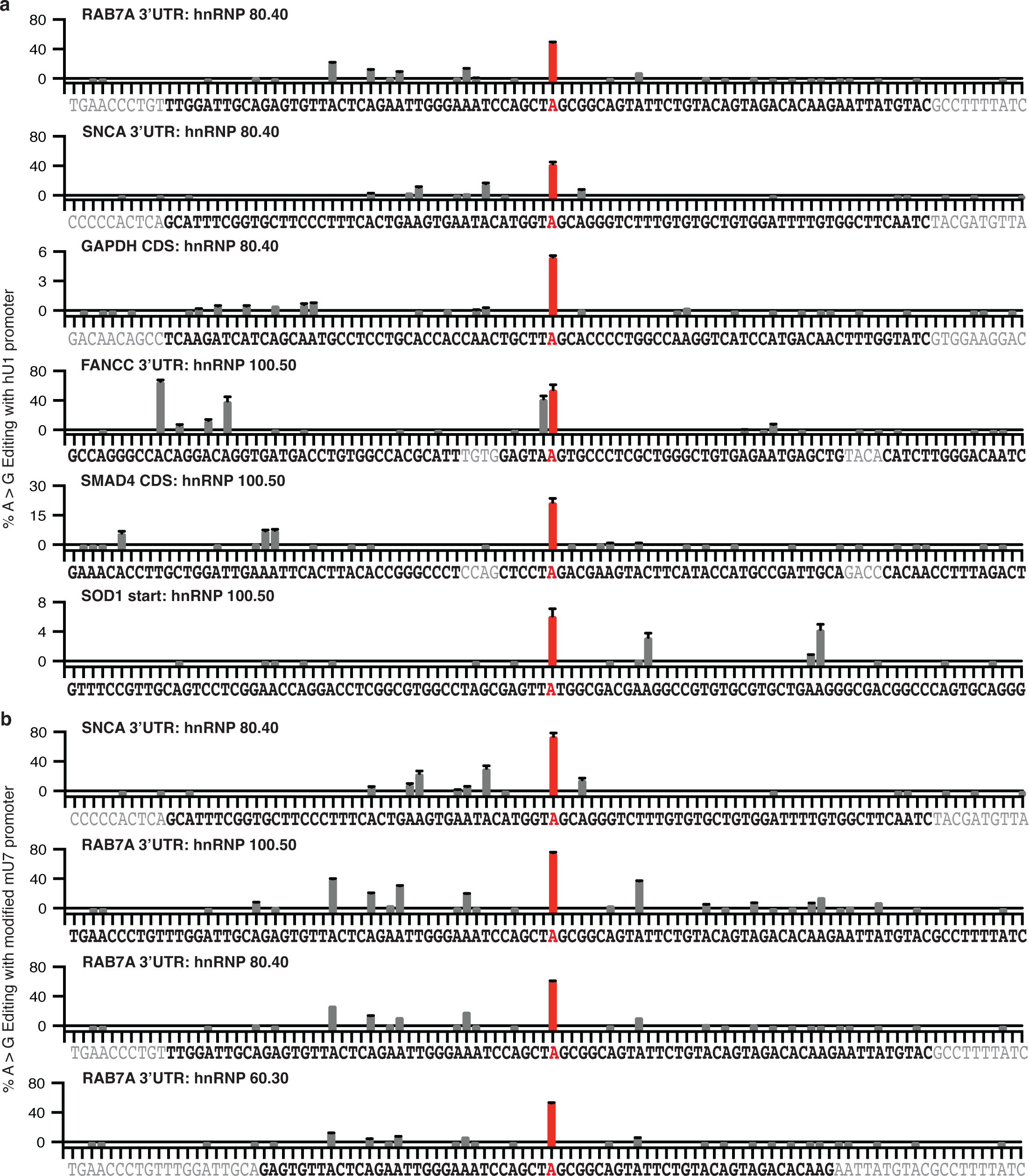
Guide RNAs with the triple variant SmOPT U7 hairpin show preferential target base editing over bystander editing. All guide RNAs possessed a 5’ double hnRNP A1 binding motif and the triple variant SmOPT mU7 hairpin sequence. Constructs were individually transfected into Landing Pad 293T cells for single copy integration; RNA was measured 13 days later. 100 nt of the target mRNA sequence is listed below; the adenosine targeted by the A-C mismatch is highlighted in red; the target sequence covered by the antisense guide RNA is in black. **a**, RNA editing profiles for six gRNA constructs expressed by the hU1 promoter in Fig. 3. **b**, RNA editing profiles for four gRNA constructs expressed by the modified mU7 promoter in Figures 4 and 5.

